# Middle-down proteomics reveals dense sites of methylation and phosphorylation in arginine-rich RNA-binding proteins

**DOI:** 10.1101/775122

**Authors:** Sean R. Kundinger, Isaac Bishof, Eric B. Dammer, Duc M. Duong, Nicholas T. Seyfried

## Abstract

Arginine (Arg)-rich RNA-binding proteins play an integral role in RNA metabolism. Post-translational modifications (PTMs) within Arg-rich domains, such as phosphorylation and methylation, regulate multiple steps in RNA metabolism. However, the identification of PTMs within Arg-rich domains with complete trypsin digestion is extremely challenging due to the high density of Arg residues within these proteins. Here, we report a middle-down proteomic approach coupled with electron transfer dissociation (ETD) mass spectrometry to map previously unknown sites of phosphorylation and methylation within the Arg-rich domains of U1-70K and structurally similar RNA-binding proteins from nuclear extracts of HEK293 cells. Remarkably, the Arg-rich domains in RNA-binding proteins are densely modified by methylation and phosphorylation compared with the remainder of the proteome, with di-methylation and phosphorylation favoring RSRS motifs. Although they favor a common motif, analysis of combinatorial PTMs within RSRS motifs indicate that phosphorylation and methylation do not often co-occur, suggesting they may functionally oppose one another. Collectively, these findings suggest that the level of PTMs within Arg-rich domains may be among the highest in the proteome, and a possible unexplored regulator of RNA metabolism. These data also serve as a resource to facilitate future mechanistic studies of the role of PTMs in RNA-binding protein structure and function.

**Briefs:** Middle-down proteomics reveals arginine-rich RNA-binding proteins contain many sites of methylation and phosphorylation.

## Introduction

Key proteins that carry out specialized biological processes such as RNA splicing, polyadenylation and transport, contain domains disproportionately enriched with arginine [1, 2]. RNA-binding proteins (RBPs) that harbor Arginine (Arg)-rich domains may be broadly classified into two subsets based on residue composition. One class of these RBPs contains highly repetitive complementary repeats of basic (K/R) and acidic (D/E) residues, that we have previously referred to as <u>B</u>asic <u>A</u>cidic <u>D</u>ipeptide (BAD) domains [3]. Importantly, BAD domains facilitate protein aggregation, and in the context of Alzheimer’s disease (AD), facilitate interactions with pathological Tau protein [3]. A second subset, related to the BAD proteins, are the Arginine/serine-rich (RS) domains that are ubiquitous in the Serine/Arginine (SR) family of proteins [2]. Upon serine phosphorylation, RS domains mimic BAD domains with a similarly alternating basic-acidic dipeptide sequence pattern. The RS domains are commonly found in splicing factors and are essential for alternative splicing, protein-protein interactions, and localization [4–9].

The functional and structural diversity of the proteome is markedly increased through post-translational modifications (PTMs), which are covalent modifications of protein primary structure, with over 300 types of PTMs identified to date [10]. The RS domains are hypothesized to be key hubs of RBP regulation, finely regulated by reversible PTM [6, 7, 11]. Serine-arginine protein kinases (SRPKs) phosphorylate RS domains to influence alternative splicing and localization of RS domain-containing proteins, indicating that the PTM status of RS domains may regulate a vast range of essential RNA metabolism processes (9-11). The phosphorylation and subsequent de-phosphorylation of Arg-rich domains controls RBP subcellular localization, spliceosome assembly and splice site selection [12, 13]. Identifying and understanding the functional consequences of PTMs in Arg-rich domains is critical to provide mechanistic insight.

Although the functional role of RS domain PTMs is evident, attempts to globally identify individual PTM sites by mass spectrometry (MS) have been stymied by technical challenges. This is largely due to the inability to sequence Arg-rich domains using bottom-up proteomic approaches. Standard bottom-up proteomic methods rely on a complete digestion of proteins using the serine protease trypsin, which cleaves at arginine and lysine residues to produce peptides generally of appropriate lengths and charge suitable for MS identification [14, 15]. While this strategy is appropriate for proteins with a normal distribution of arginine and lysine, it is ineffective when examining proteins with Arg-rich domains. Complete digestion of Arg-rich domains using trypsin produces peptides that are too small to map to a unique protein. Additionally, peptides produced from Arg-rich domains are highly protonated and thus difficult to fragment by standard approaches. Peptide fragmentation by tandem mass spectrometry with collision induced dissociation (CID) or higher-energy collision dissociation (HCD) disfavors multiply-protonated Arg-rich peptides [16–18]. Arginine, the most basic amino acid, contains three basic nitrogens that collectively reduce proton mobility along the peptide backbone, eventually sequestering protons and preventing dissociation [19, 20].

Electron-transfer dissociation (ETD) is a non-ergodic fragmentation technique, related to electron capture dissociation (ECD), that cleaves the peptide backbone N-C_α_ bond following the transfer of an electron from an anion (e.g., fluoranthene radical ion) of low electron affinity to a multiply-charged peptide [17, 18, 21, 22]. This dissociation event results in c-and z-type fragment ions instead of the typical b-and y-type ions observed in CID or HCD [23, 24]. Notably, ETD peptide fragmentation is not influenced by amino acid side-chain chemistry and thereby preserves PTMs that are otherwise labile by CID, such as phosphorylation, glycosylation, nitrosylation and sulfonation, providing more complete PTM identification [16, 21, 22, 25–32]. ETD has been extensively utilized to map PTMs within highly basic N-terminal histone tails and similar high charge state peptides by both bottom-up and middle-down approaches [30–33]. Non-canonical ‘middle-down’ approaches may be leveraged to characterize longer peptides by employing shorter proteolysis time points or non-canonical proteolytic enzymes. This is in contrast to standard ‘bottom-up’ strategies, that utilize overnight trypsin digestion, typically rendering Arg-rich domains extremely difficult to detect by mass spectrometry. Importantly, the middle-down approaches enhanced the ability to both sequence Arg-rich protein sequences and capture co-occurring or combinatorial PTMs [30, 31].

Here we report a middle-down proteomic approach utilizing limited trypsin digestion and ETD on an Orbitrap Fusion mass spectrometer to map PTMs within Arg-rich domains [34]. We achieved near complete coverage and identification of multiple phosphorylation and methylation sites within the Arg-rich domains from the core snRNP protein U1 small nuclear ribonucleoprotein 70 kDa (U1-70K), among other key splicing RBPs within the nucleoplasm of HEK293 cells. We found BAD and RS domains alike to be decorated with methylation of arginine and lysine residues and phosphorylation of serine, threonine and tyrosine residues. Peptides mapping within these domains contained a combinatorial pattern of PTM, enriched above background proteomic levels. These data suggest that Arg-rich domains are highly dense sites of PTM, and that PTMs could play a critical role in maintaining protein-protein interactions. This approach can be generally applied for mapping PTMs in arginine- or lysine-rich domains found throughout the proteome.

## Materials and Methods

### Materials

The following are primary antibodies included in this study: an in-house rabbit polyclonal antibody raised against a synthetic keyhole limpet hemocyanin-conjugated peptide corresponding to an epitope C-terminal to the low complexity domains of U1-70K [35]; anti-SRSF1 antibody (catalog no. ab38017, Abcam). Secondary antibodies were conjugated to either Alexa Fluor 680 (Invitrogen) or IRDye800 (Rockland) fluorophores.

### Plasmids and Cloning

The U1-70K LC1/BAD plasmid used herein was generated in a previous study [35]. Cloning was performed by the Emory Custom Cloning Core Facility (Oskar Laur), and plasmids were confirmed by DNA sequencing.

### Cell Culture and Transfection

Human embryonic kidney (HEK) 293T cells (ATCC CRL-3216) were cultured in Dulbecco’s Modified Eagle Medium (DMEM, high glucose (Gibco)) supplemented with 10% (v/v) fetal bovine serum (Gibco) and 1% penicillin-streptomycin (Gibco), and maintained at 37°C under a humidified atmosphere of 5% (v/v) CO_2_ in air. For transient transfection, the cells were grown to 80-90% confluency in 10 cm^2^ culture dishes and transfected with 10 μg expression plasmid and 30 μg linear polyethylenimine (PEI).

### Protein expression and purification

Recombinant U1-70K protein was expressed and purified as described previously [36]. In short, HEK293T cells were grown and transfected with a plasmid encoding the GST-LC1/BAD (AA231-310) domain of U1-70K using PEI, and harvested 72 hours post-transfection. The cells were resuspended in ice-cold lysis buffer (50 mM HEPES pH 7.4, 200 mM NaCl, 5% glycerol, 1 mM EDTA, 1% (w/v) sarkosyl and 1X HALT protease/phosphatase inhibitor cocktail). Lysates were sonicated using a microtip probe to shear nucleic acids. To make the lysates compatible with GST affinity purification, Triton X-100 (TX-100) was added to a concentration of 1.5% (v/v). The lysates were cleared by centrifugation at 14,000 x g for 10 min at 4°C. The resulting supernatants were incubated overnight at 4°C with 0.5-1 ml of swelled glutathione-agarose resin (Sigma G4510), after which the slurries were loaded onto a column. The resin was washed with 10 column volumes of wash buffer (50 mM HEPES pH 7.4, 200 mM NaCl and 1% TX-100), and eluted with 4 x 1 ml elution buffer (50 mM Tris-HCl pH 8.5, 500 mM NaCl, 20 mM reduced L-glutathione (Sigma G4251) and 0.1% TX-100. The eluted fractions were concentrated to ∼200 μl using Amicon Ultra-0.5 ml 10K MWCO Centrifugal Filter Units (EMD Millipore), and dialyzed overnight against 50 mM HEPES pH 7.4, 200 mM NaCl and 0.1 mM PMSF using 10K MWCO Slide-A-Lyzer MINI Dialysis Units (Thermo). Protein concentration was determined by running each elution fraction on an SDS-PAGE gel with bovine serum albumin (BSA) standards ranging from 0.2-1 μg per lane and staining with Coomassie G-250 [36]. Densitometry of the BSA standards was used to calculate the concentration of GST affinity purified protein.

### Western Blotting

Western Blotting was performed according to standard protocol as previously described in Bishof *et al.* [3]. In short, samples were boiled in Laemmli sample buffer (8% glycerol, 2% SDS, 50mM Tris pH 6.8, 3.25% beta-mercaptoethanol) for 5 minutes, then resolved on a Bolt® 4-12% Bis-tris gel (catalog no. NW04120BOX, Invitrogen) by SDS-PAGE and semi-dry transferred to a PVDF membrane with the iBlot2 system (ThermoFisher). Membranes were blocked with TBS Starting Block Blocking Buffer (catalog no. 37542, ThermoFisher) and probed with primary antibodies (1:1,000 dilutions) overnight at 4°C. Membranes were then incubated with secondary antibodies conjugated to either Alexa Fluor 680 (Invitrogen) or IRDye800 (Rockland) fluorophores for one hour at RT. Membranes were imaged using an Odyssey Infrared Imaging System (Li-Cor Biosciences) and band intensities were calculated using Odyssey imaging software.

### In-gel Limited Trypsin digestion

The GST-LC1/BAD (residues 231-310) purified protein (14 µg) was run onto a 10% acrylamide gel. The gel was stained with Coomassie G-250 and the GST-LC1/BAD band was compared to BSA standards to estimate protein concentration. The GST-LC1/BAD band was cut out and diced into small pieces. The gel pieces were divided among five tubes each receiving ∼3 µg of GST-LC1/BAD. Gel pieces were de-stained until clear using 70% 50mM ammonium bicarbonate (ABC) and 30% acetonitrile. While on ice, each tube received 30µL digest buffer [12.5 ng/µl trypsin (Pierce MS grade) in 50mM ABC buffer. After the addition of trypsin, samples were incubated on ice for 3 minutes then brought up to room temperature to start digestion. At 6 different time points (15, 30, 60, 120, 240 minutes, and overnight) excess trypsin solution was removed and the digestion reaction was stopped with 30 µl extraction buffer (50% acetonitrile, 5% acetic acid). After the addition of extraction buffer, samples were allowed to equilibrate for five minutes, then stored at −20 °C until peptide extraction. Peptides were shaken for 40 minutes at room temperature then spun at 20,000 x g for one minute. After a minute period following the first spin, peptides were spun again. This spin and relax cycle was repeated twice. The supernatant containing the extracted peptides was then collected into a new tube, in which 30 µl of extraction buffer was then added. This extraction process was repeated two more times. Peptides were lyophilized using a SpeedVac (catalog no. 731022, Labconco) and resuspended in MS sample loading buffer (1% acetonitrile, 0.1% formic acid, and 0.03% trifluoroacetic acid).

### Nucleoplasm Enrichment

This cellular extraction procedure was adapted from the Gozani group [37]. In short, cells from eight 15-cm plates were combined and rinsed with cold PBS, and then scraped in 10 ml PBS + 1X Protein Inhibitor Cocktail buffer (catalog no. COUL-RO, Roche) and centrifuged for 5 min at 1,000 x g at 4°C. Cells were washed once in 1 ml 1X (cold) PBS and recovered by centrifugation for 5 min at 1,000 x g at 4°C, swelled in 75 ul of hypotonic lysis buffer (10 mM HEPES pH 7.9, 20 mM KCl, 0.1 mM EDTA, 1mM DTT, 5% Glycerol, 0.5 mM PMSF, 10 ug/ml Aprotinin, 10 ug/mL Leupeptin) and incubated on ice for 10 min. Samples were lysed by 0.1% NP-40, vortexed, and incubated on ice for 5 min. Nuclei were recovered by centrifugation for 10 min at 15,600 x g at 4°C. Nuclei were extracted for 30 min on ice in 40 ul high salt buffer (20 mM HEPES pH 7.9, 0.4 M NaCl, 1 mM EDTA, 1 mM EGTA, 1 mM DTT, 0.5 mM PMSF, 10 ug/ml Aprotinin, 10 ug/mL Leupeptin). Samples were sonicated for 5 sec and extracts were collected by centrifugation for 10 min at 15,600 x g at 4°C. The supernatant obtained following the centrifugation step consisted of isolated nucleoplasm and the resulting pellet was the chromatin fraction.

### In-solution digest of nucleoplasm fractions

For each time point, 150 µg of nucleoplasmic fraction was digested. Samples were brought to a final concentration of 1M urea, then dithiothreitol was added to a final concentration of 1 mM, and incubated for 30 minutes. Iodoacetamide was added to a final concentration of 1 mM, and incubated for 20 minutes in the absence of light. Samples were then diluted in digestion buffer, and digested with a 1:50 ratio of trypsin (Pierce MS grade) to total protein. The digestion reactions were performed at room temperature and quenched at increasing time lengths (5, 10, 20, 40, 80, 160 minutes, overnight) with 0.1% formic acid, and 0.01% trifluoroacetic acid solution. Resulting peptides were cleaned up using an HLB column (Waters). Samples were washed first with methanol, then Buffer C (50% acetonitrile and 50% water), then 0.1% trifluoroacetic acid. The digested samples were then loaded into the column, washed with 0.1% trifluoroacetic acid twice, and then eluted with Buffer C. The resulting elutant was lyophilized using a SpeedVac (catalog no. 731022, Labconco).

### Mass spectrometry analysis

Lyophilized peptides were resuspended in loading buffer (0.1% formic acid, 0.03% TFA, 1% acetonitrile) and separated on a self-packed C18 (1.9 μm Dr. Maisch, Germany) fused silica column (20 cm × 75 μm internal diameter; New Objective, Woburn, MA) by a NanoAcquity UHPLC (Waters). Linear gradient elution was performed using Buffer A (0.1% formic acid, 0% acetonitrile) and Buffer B (0.1% formic acid, 80% acetonitrile) starting from 3% Buffer B to 40% over 100 min at a flow rate of 300 nl/min. Mass spectrometry was performed on an Orbitrap Fusion Tribrid Mass Spectrometer. Data-dependent MS/MS analyses included a high resolution step (120,000 at m/z 400) with an m/z range of 100-1000. MS1 scans were conducted in the Orbitrap, and the top 10 ions with highest charge, followed by the precursor ion with the greatest intensity, were given priority for fragmentation. A data-dependent decision tree was used [38] and MS/MS spectra from both HCD and ETD were collected in the ion-trap. Peptides with charge state of +2 were chosen for fragmentation by HCD only, while all charge states +3 and above were fragmented by both ETD and HCD. At +3 charge state, precursor ions under 650 m/z were fragmented by both ETD and HCD. Those equal to or greater than 650 m/z were fragmented by HCD only. At +4 charge state, precursor ions under 900 m/z were jointly fragmented by both HCD and ETD, and only by HCD at m/z values equal to greater than 900. At +5 charge state, precursor ions under 950 m/z were fragmented by both HCD and ETD, and only by HCD at m/z values equal to or greater than 950. At + 6 charge state, all precursor ions were fragmented by ETD. Dynamic exclusion was set to exclude previous sequenced precursor ions for 30 seconds.

### Database Searching

Data files for the time points were analyzed using MaxQuant v1.5.2.8 and v1.6.1.0 with Thermo Foundation 2.0 for RAW file reading capability. The search engine Andromeda was used to build and search a concatenated target-decoy UniProt Knowledgebase (UniProtKB) containing both Swiss-Prot and TrEMBL human reference protein sequences (90,411 target sequences downloaded April 21, 2015), plus 245 contaminant proteins included as a parameter for Andromeda search within MaxQuant [39]. Methionine oxidation (+15.995 Da), and protein N-terminal acetylation (+42.011 Da) were included as variable modifications (up to 5 allowed per peptide); cysteine was assigned a fixed carbamidomethyl modification (+57.022 Da) for the nucleoplasm. A two stage search was performed as described previously [40]. This two-step method allows for a larger search space while limiting false-discovery rate (FDR) [41]. For the first search only fully tryptic peptides were considered with up to 2 missed cleavages in the database search. A precursor mass tolerance of ± 20 ppm was applied prior to mass accuracy calibration and ± 4.5 ppm after internal MaxQuant calibration. Other search settings included a maximum peptide mass of 6,000 Da, a minimum peptide length of 6 residues and 0.6 Da tolerance for ion-trap ETD and HCD MS/MS scans. The false discovery rate (FDR) for peptide spectral matches, proteins, and site decoy fraction were all set to 1 percent. The proteins identified in the first database search were then used to make a targeted FASTA formatted database to re-search the raw files with an expanded missed cleavage window and variable modifications. The targeted database corresponding to purified U1-70K LC1/BAD protein contained a FASTA file of 366 proteins and allowed for 6 mis-cleavages. The targeted database corresponding to the nucleoplasm contained a FASTA file of 4,307 unique protein groups (20,392 protein isoforms), allowing 6 mis-cleavages. Both targeted databases were used to search the spectra for variable PTMs including phosphorylation (S/T/Y) (+79.966 Da) and mono- and di-methylation (K/R) (+14.016 Da, +28.031 Da), in addition to N-terminal acylation and methionine oxidation. The MS/MS spectra were collected by a low-resolution ion trap (product ion tolerance = 0.6 Da). As such, trimethylation of lysine (42.047 ± 0.002 Da) and acetylation of lysine (42.011 ± 0.004 Da) were not included in our search, given that the masses of these modifications are less than 20 ppm apart. The .raw and .txt files obtained from MaxQuant searches were uploaded to ProteomeXchange on 9/4/2019 (Accession ID: PXD015208).

### Parameters for selecting Basic Acidic Dipeptide (BAD) and Arginine-Serine (RS) peptides

We applied a BAD score algorithm counting peptides that added 1.0 points for an alternating basic and acidic charge (+/- or -/+) and added 0.1057/0.0764/0.0475 points for S/T/Y residues, respectively, neighboring a basic residue. This attempted to reflect the average relative phosphorylation frequency across all nucleoplasmic peptides of each residue, which transforms the dipeptide into a BAD sequence. This sum was calculated and divided by the peptide length to give a “BAD score” for each peptide. A cutoff score of 0.3 or greater was chosen, identifying 543 BAD peptides in total. These selection criteria enriched for Lys/Arg/Ser/Thr/Tyr residues, and actual residue frequency was factored into normalized PTM fold changes. RS peptides were identified through a similar algorithm, counting peptides with 1.0 points for alternating Arg-Ser/Ser-Arg residues, normalized to peptide length. The sum was calculated and divided by the peptide length to give an “RS score” for each peptide. A cutoff of greater or equal to 0.2 was chosen. There were 533 RS peptides identified by this selection algorithm. These selection criteria resulted in an enrichment for Lys/Arg/Ser/Thr/Tyr residues, which was factored into normalizing PTM fold changes for RS peptides.

### MEME analysis

Phosphosite Motif Logo analysis was conducted (https://www.phosphosite.org/sequenceLogoAction.action) [42] to identify sequence motifs of serine phosphorylation, arginine monomethylation and arginine dimethylation. Input peptide sequence windows with unique sites of PTM were assembled and entered for analysis. A list of 46 monomethylated arginine peptides, 252 dimethylated arginine peptides and 458 phosphorylated serine peptides generated by our MS data served as input sequences. Additionally, Motif analysis was performed for mono-/di-methylated lysines, and threonine/tyrosine phosphorylation (Supplemental Fig. S5).

### Sequence Coverage Code

Peptide Coverage Summarizer software v1.3.6794.31818 (Pacific Northwest National Laboratory, Richland, WA) was used to input unique peptide sequences of all peptide spectral matches identified in the middle-down ETD LC-MS/MS run and search results of this study’s MaxQuant re-search, with up to 6 miscleavages. Coverage was summarized by the software as capitalized single-letter encoded residues in output of the full sequence of each protein in the UniprotKB human FASTA database (90,411 proteins). An in-house R script compared these residues with coverage summarized against the same database obtained by the Dec 2018 Proteome Atlas (1,817,080 unique peptides; downloaded February 26, 2019). Then residues uniquely matched by our search and not by previous efforts were identified by comparison of the two coverage summaries and tallied. Degenerate peptides matching to several proteins were not included. The longest isoform was chosen for each gene product’s set of protein isoforms, when at least one isoform contained novel coverage relative to Proteome Atlas. Sublists of BAD and RS score threshold-passing peptide sequences identified in the same experiment were inputted separately through the pipeline to tabulate novel coverage statistics for these two special classes of peptide.

### Measurement of PTM Coverage and Motif Occupancy

Modified sequences containing a 31-residue width window were matched to sequences downloaded (5/24/2019) from phosphosite.org [42]. Excel was used to match methylated or phosphorylated sequences previously observed by mass spectrometry methods only. A character length (len) function in Excel substitution function was used to search for post-translationally modified RSRS/SRSR sequences. Occurrences were normalized to the number of RSRS/SRSR sequences within a peptide and calculated as percent of total occurrences.

## Results

As shown in Fig. 1A, the spliceosomal protein, U1-70K, contains two Arg-rich low complexity (LC) domains, LC1/BAD (residues 231-310) and LC2 (residues 317-407). The ‘BAD’ acronym of LC1/BAD stands for “<u>B</u>asic <u>A</u>cidic <u>D</u>ipeptide”, containing dipeptide repeats of a basic (K/R) residue adjacent to an acidic residue (D/E). The U1-70K LC1/BAD domain has both BAD and RS motifs. The LC1/BAD domain of U1-70K is of particular biological interest due to its central role in U1-70K nuclear localization, granule formation, and co-aggregation with Tau in Alzheimer’s disease [3, 35, 43]. There are currently over 50,000 peptide spectral matches to U1-70K using standard bottom-up approaches with complete trypsin digestion, mapping approximately 70% of the protein [44]. Despite this number of spectral matches, only 7.5% of the LC1/BAD domain has previously been sequenced, or 6 out of 80 residues [45, 46] (Fig.1A). Thus, we sought to develop a method to sequence the U1-70K LC1/BAD domain that would be generally applicable to other Arg-rich RNA-binding proteins in the proteome.

**Figure 1.**
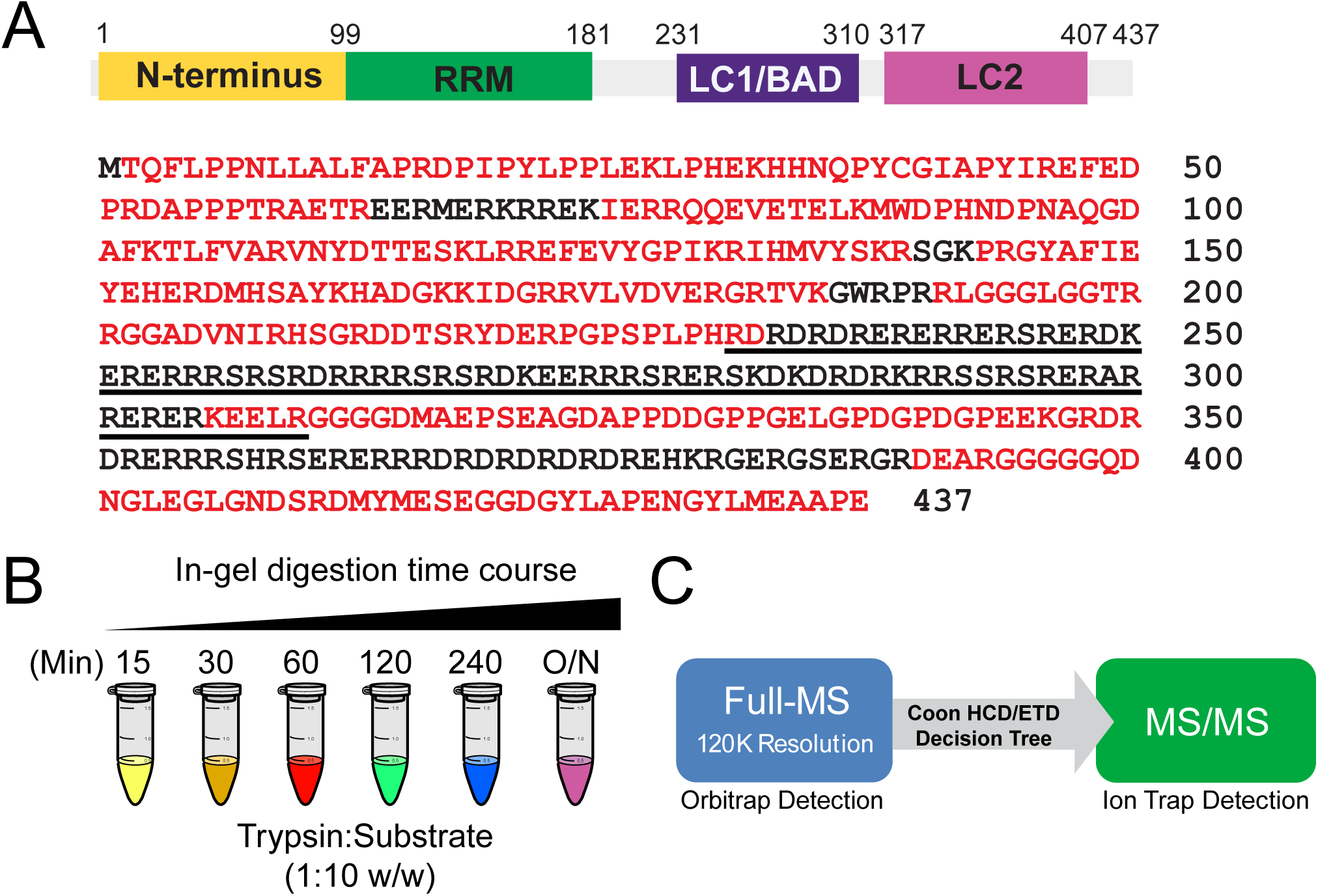
Middle-down Electron Transfer Dissociation (ETD) Mass Spectrometry (MS) strategy. ***(A)*** Schematic of U1-70K protein domains. Proteomic coverage map of U1-70K with residues previously observed by mass spectrometry and uploaded to peptideatlas.org (*red*) and residues that lack mapped peptides (*black*). The LC1/BAD domain (*underlined*) is almost empty of MS coverage. ***(B)*** At six different time points [15, 30, 60, 120, 240 minutes, and overnight (O/N)] the digestion reaction was quenched with acetic acid. ***(C)*** Extracted peptides were analyzed by LC-MS/MS on an Orbitrap Fusion using a data-dependent decision tree acquisition method [28] to alternatively select between HCD and ETD peptide fragmentation based on the charge state and m/z of the precursor ion. A two-step database search method using the Andromeda search engine was employed wherein proteins that were identified from a primary target-decoy search against human UniProtKB database were used to create a second smaller focused database. This focused database was then used to search for phosphorylation (serine, threonine or tyrosine) and mono-, di- and tri-methylation (arginine or lysine) with consideration of up to 6 missed trypsin cleavage events.

A combination of electron transfer dissociation (ETD) and limited trypsin proteolysis (middle-down) strategies were used to sequence the LC1/BAD domain of U1-70K. Recombinant LC1/BAD domain was transiently expressed and GST-purified from human embryonic kidney (HEK293T) cells. The LC1/BAD protein (∼38 kDa) was incubated with trypsin for varying time periods to achieve a partial digest. The digestion reaction was quenched with acetic acid at 15, 30, 60, 120, and 240 minutes. An additional complete digestion sample was also allowed to proceed overnight (Fig.1B). The resulting peptides were then extracted and analyzed by LC-MS/MS on an Orbitrap Fusion mass spectrometer. Higher-energy collisional dissociation (HCD) and electron transfer dissociation (ETD) decision tree was used (Fig.1C) [28].

In order to sequence low complexity Arg-rich domains, we needed to account for a large number of missed cleavages and PTMs, which would increase the search space compared to traditional proteome database searches. A two-step database search method was utilized wherein proteins matched from a primary search were used to create a smaller, focused database. This strategy limits the false discovery rate (FDR) and the number of false negatives [41], resulting in higher confidence peptide spectral matches (PSMs) compared to traditional one-step database search methods [41]. Our focused database was then used to perform a second search that included increased missed-cleavages, and PTMs (methylation and phosphorylation), using a <1% FDR cutoff.

A strength of ETD is the ability to fragment highly protonated peptides and retain PTMs [16, 25, 27, 28, 30–32]. For example, when comparing the MS/MS spectra of the same peptide fragmented by either ETD or HCD, the ETD method results in increased fragmentation (Fig.2A-B). These fragment ions (c- and z-ions) produced by ETD provide unique diagnostic ions, unafforded by canonical HCD/CID fragmentation methods, that enhance identification of the peptide (Fig.2B) [47, 48]. When examining the HCD spectra, only a neutral loss of 98 Da is observed. This mass shift is a signature of a phosphorylation PTM, indicating the loss of phosphate and water [49–52] (Fig.2A). However, the HCD spectra contain few other additional fragment ions (b- and y-ions), making it difficult to map the linear sequence of amino acids of the precursor peptide. When observing unique peptide matches accumulated across all time points, as a result of increased fragmentation, ETD yielded 114 unique LC1/BAD peptides, as compared to HCD, which identified only 18 unique LC1/BAD peptides (Fig.2C).

**Figure 2.**
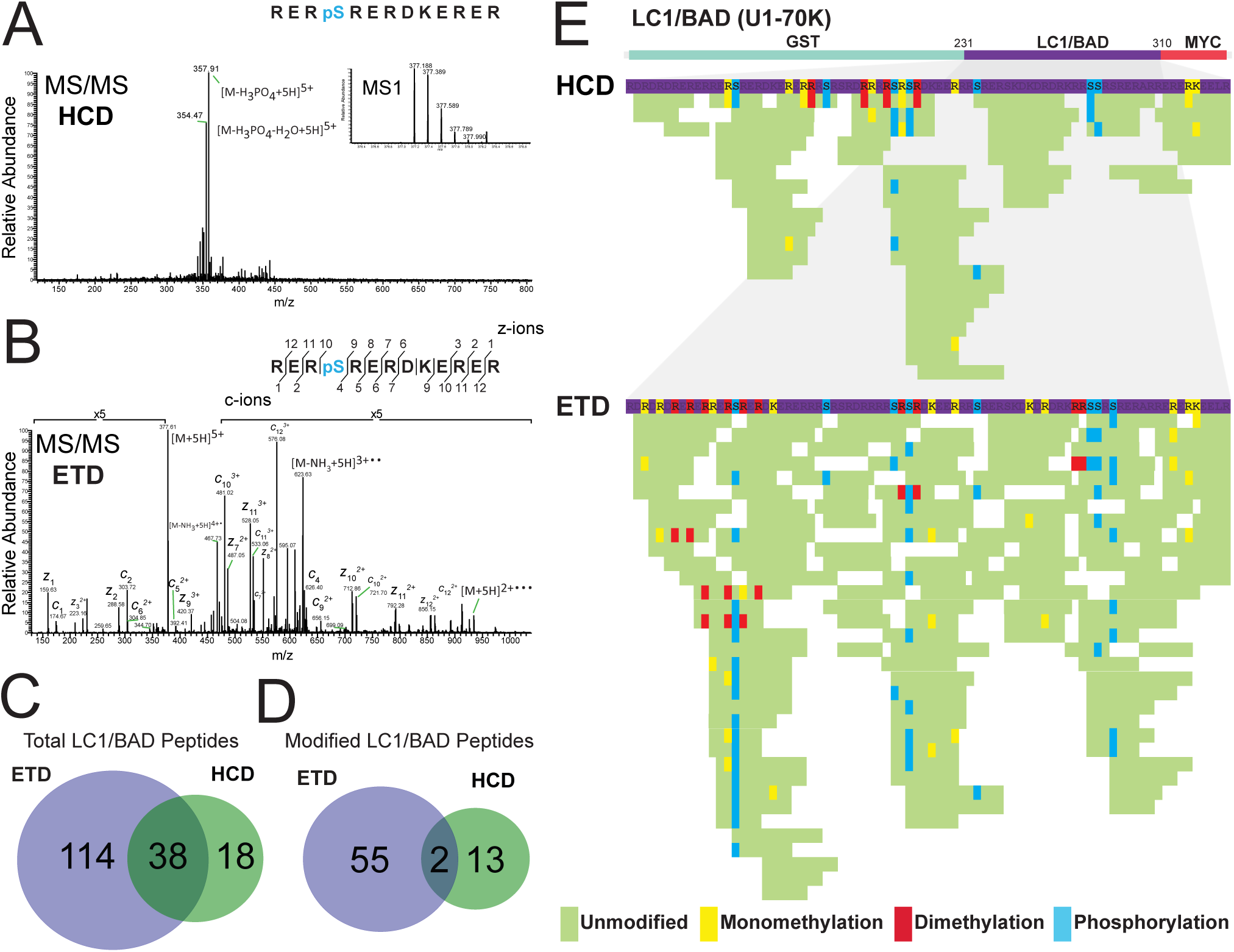
ETD more effectively fragments Arg-rich peptides compared to HCD. *(**A**)* HCD MS/MS output of a R242-R254 U1-70K LC1/BAD peptide. HCD results in a strong neutral loss (−98 Da) of phosphoric acid and few fragment ions (b- and y-ions). *(**B**)* Isolation and ETD fragmentation of a charged-reduced precursor isotope [M-NH_3_+5H]^3+•••^ of the same peptide reveals an information-rich MS/MS spectrum (c- and z-ions), while preserving the phosphorylated serine, labile by HCD. ***(C)*** ETD fragmentation produces the vast majority of unique peptide matches to U1-70K LC1/BAD as compared to HCD. ***(D)*** ETD fragmentation facilitates the identification of increased number of post-translationally modified peptides matched to U1-70K LC1/BAD as compared to HCD. ***(E)*** Protein coverage generated by HCD fragmentation is compared ETD fragmentation across all trypsin incubation lengths (15, 30, 60, 120, 240 minutes, and overnight). Peptides are colored according to whether each residue is unmodified (*green*), mono-methylated at an Arginine/Lysine (*yellow*), di-methylated at Arginine/Lysine (*red*) or phosphorylated at Serine/Threonine/Tyrosine (*turquoise*). ETD outperforms HCD in preserving PTMs and sequencing highly-protonated regions of recombinant U1-70K LC1/BAD domain.

When examining LC1/BAD peptides containing PTMs, the difference between peptides identified by ETD and HCD fragmentation is similarly apparent. ETD identifies 55 PTM-containing LC1/BAD peptides alone compared with HCD, which identified just 13 such peptides (Fig.2D). To visualize these findings, the LC1/BAD unique peptides identified by either HCD or ETD fragmentation were mapped onto the LC1/BAD domain (Fig.2E). ETD led to the identification of more LC1/BAD unique peptides than HCD, providing coverage of 79/80 residues (99%) within the LC1/BAD domain (**Supplemental Table S1)**. If trypsin digestion is allowed to proceed overnight, the number of PSMs decreases and significant coverage is no longer obtained with either ETD or HCD. Using the middle-down ETD strategy, we achieved near complete coverage of the LC1/BAD domain of U1-70K *in vitro*.

Thus, the combination of both ETD and a limited digest is necessary for complete coverage of the LC1/BAD domain of U1-70K. Furthermore, we identified 36 sites of PTMs summed across all proteolysis time lengths (Supplemental Fig. S4A). Eight out of 10 possible serine residues within LC1/BAD were found to be phosphorylated. Additionally, 23 total sites of arginine methylation were discovered, as 12 of these arginines were mono-methylated while 13 arginine residues were dimethylated. Furthermore, four sites of lysine mono-methylation were discovered. No lysine di-methylation was observed in the LC1/BAD domain of recombinant U1-70K LC1/BAD protein. This near complete coverage of purified recombinant U1-70K LC1/BAD serves as proof of concept for using a middle-down ETD to sequence Arg-rich proteins in complex mixtures.

### Preparation of nucleoplasm fractions enriched with Arg-rich RBPs

We next sought to employ this approach globally to achieve widespread coverage of Arg-rich BAD and RS proteins, mapping PTMs therein, from complex cell extracts. Arginine is not evenly distributed throughout the proteome, but rather is densely concentrating in Arg-rich domains [1, 2]. Small nuclear ribonucleoproteins (snRNPs), associated heterogeneous nuclear ribonucleoproteins (hnRNPs) and serine/arginine-rich splicing factors (SRSFs) form macromolecular spliceosome structures in the nucleus, many of which contain Arg-rich domains [53, 54]. To obtain a biological sample enriched in RBPs with Arg-rich domains, cellular fractionation was performed to isolate the nucleoplasm. The nucleoplasm is rich in splicing RBPs, many of which undergo liquid-liquid phase separation (LLPS) and aggregate in neurodegenerative disease [3, 36, 55, 56]. For downstream proteomic analysis, the over-representation of highly-expressed histone proteins in nuclear fractions would suppress the sensitivity of the mass spectrometer to identify comparatively less abundant RBPs that we sought to capture in our analyses. Therefore, simultaneous nucleoplasm isolation and histone depletion was performed to increase the probability of sequencing Arg-rich RBPs, especially low stoichiometry peptides within BAD and RS domains, many of which are expected to contain PTMs.

Differential centrifugation was performed to enrich cytoplasmic, nuclei, chromatin and nucleoplasmic fractions from HEK-293 cell extract (Fig.3A). Following nucleoplasm isolation, core splicing proteins U1-70K and SRSF1 were analyzed by SDS-PAGE and western blotting, showing a 2.75- and 7.02-fold enrichment in the nucleoplasmic fraction, respectively, relative to the original whole cell fraction (Fig.3B-C). Importantly, coomassie staining of an identical gel showed histone proteins were enriched by 5.26-fold in the chromatin fraction (Fig.3B-C), and depleted from the nucleoplasm by 0.45-fold, relative to the whole cell fraction. The enrichment of splicing factors and other RBPs make this complex nucleoplasm sample ideal for examination of the Arg-rich proteome.

**Figure 3.**
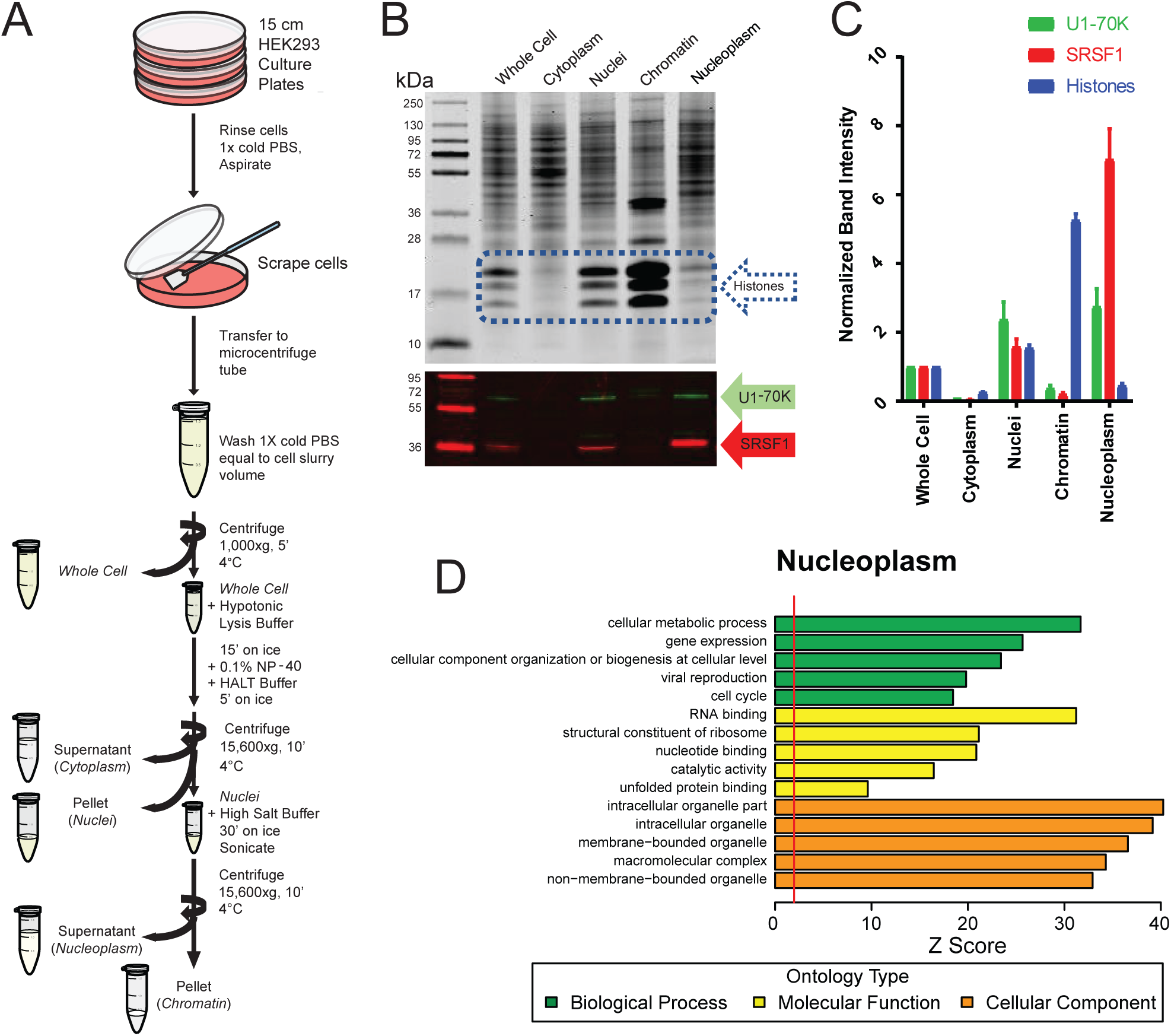
Isolation of nucleoplasm by cellular fractionation enriches for RNA-binding proteins. ***(A)*** Schematic of the HEK293T cellular fractionation protocol. ***(B)*** At the top of the panel, the individual fractions were resolved by SDS-PAGE and stained with Coomassie Blue to visualize protein. Histone proteins at the indicated molecular weight span approximately 15-22 kDa and are outlined (*dashed blue box*). At the bottom of the panel, immunoblot analysis for U1-70K (*green*) and SRSF1 (*red*) in cellular fractions. ***(C)*** Densitometry quantification of nuclear histone and RBP proteins across cellular fractions. Histone proteins show a 5.26-fold enrichment in the chromatin fraction, and 0.45-fold depletion in the nuclear extract fraction relative to the whole cell homogenate. U1-70K shows a 2.75-fold enrichment in the nucleoplasm as compared to levels in the whole cell homogenate. Similarly, SRSF1 shows a 7.02-fold enrichment in the nucleoplasm compared to whole cell homogenate. ***(D)*** Gene ontology (GO) analysis of nucleoplasm proteins identified by middle-down proteomics. Significant over-representation of an ontology term within the nucleoplasm sample compared to the background proteome is reflected with Z score greater than 1.96, equivalent to p<0.05 (to the right of the red line).

### Global analysis of Arg-rich RNA-binding from nucleoplasm extracts by middle-down ETD MS

Nucleoplasm samples were incubated with trypsin, and reactions were quenched with acetic acid at 5, 10, 20, 40, 80, and 160 minutes, with the standard overnight digestion serving as a control. Partially-digested peptides were extracted and analyzed by LC-MS/MS on an Orbitrap Fusion Tribrid mass spectrometer operating on an HCD-ETD decision tree as described for the recombinant U1-70K LC1/BAD domain expressed in HEK293T cells [28]. An initial conventional search identified 49,307 unique peptides, matching to 20,392 proteins, collapsed into 4,307 unique protein groups by parsimony. A second search using a smaller targeted database containing the 20,392 proteins matched in the first search was performed with mono-/ di-methylation of Arginine/Lysine and phosphorylation of Serine/Threonine/Tyrosine PTMs and a maximum of 6 missed cleavages. A total of 61,283 unique peptides were identified following a target-decoy search, using an FDR cutoff at less than 1%. Detected peptides matched to 4,140 unique protein groups (**Supplemental Table S2**). Although fewer input protein database entries were included in the second search, it yielded significantly more matched unique peptides (∼12,000) compared to the conventional database searched consistent with previous two-step search strategies [40]. Peptides with three or more missed-cleavages (n=8,398) and modified peptides (n=7,201) made up a significant proportion of the novel peptide matches in the two-step search strategy. Furthermore, our second database search matched 96% of protein groups identified in the first search (4,140/4,307). GO-term analysis of these proteins sequenced demonstrate that the nucleoplasm fraction was enriched with factors involved in ‘RNA binding’, consistent with our western blot results (Fig.3D). Thus, the cellular fractionation method successfully isolated nucleoplasmic proteins, enriching for native BAD and SR RBPs while also depleting histones, generating a complex sample amenable to middle-down ETD analysis.

### Enhanced sequencing coverage of Arg-rich proteins by middle-down ETD MS

We attempted to compare the influence of divergent HCD or ETD fragmentation strategies on sequencing events across the complex nucleoplasm sample. Notably, ETD and HCD sequenced a similar number of unique peptides with relatively little overlap. A total of 23,398 and 29,891 unique peptides were sequenced by ETD and HCD, respectively, while 8,239 peptides were identified by both fragmentation methods (Fig.4A). Peptides above +2 charge are preferentially sequenced by ETD as directed by the ETD/HCD decision tree algorithm [28], contributing to this peptide sequencing disparity (Supplemental Fig. S1C) This indicates that for complex sample mixtures, a decision tree approach that utilizes both ETD and HCD fragmentation platforms may provide complementary sequence information than currently achieved with standard sample preparation methods [16, 23–26]. For example, at shorter trypsin proteolysis times, ETD fragmented-peptides consisted the majority of sequenced peptides (Supplemental Fig. S1A). Between 20-40 minutes of digestion, however, HCD fragmentation becomes the preferred method of fragmentation, contributing the majority of matched peptides from 40 minutes of digestion on. Importantly, peptides sequenced following ETD fragmentation were generally more confidently scored and assigned a peptide sequence, averaging a higher Andromeda Score (147.86), a measure of the confidence of the peptide sequence assignment, as compared to HCD-fragmented peptides (103.31) (Fig.4B).

**Figure 4.**
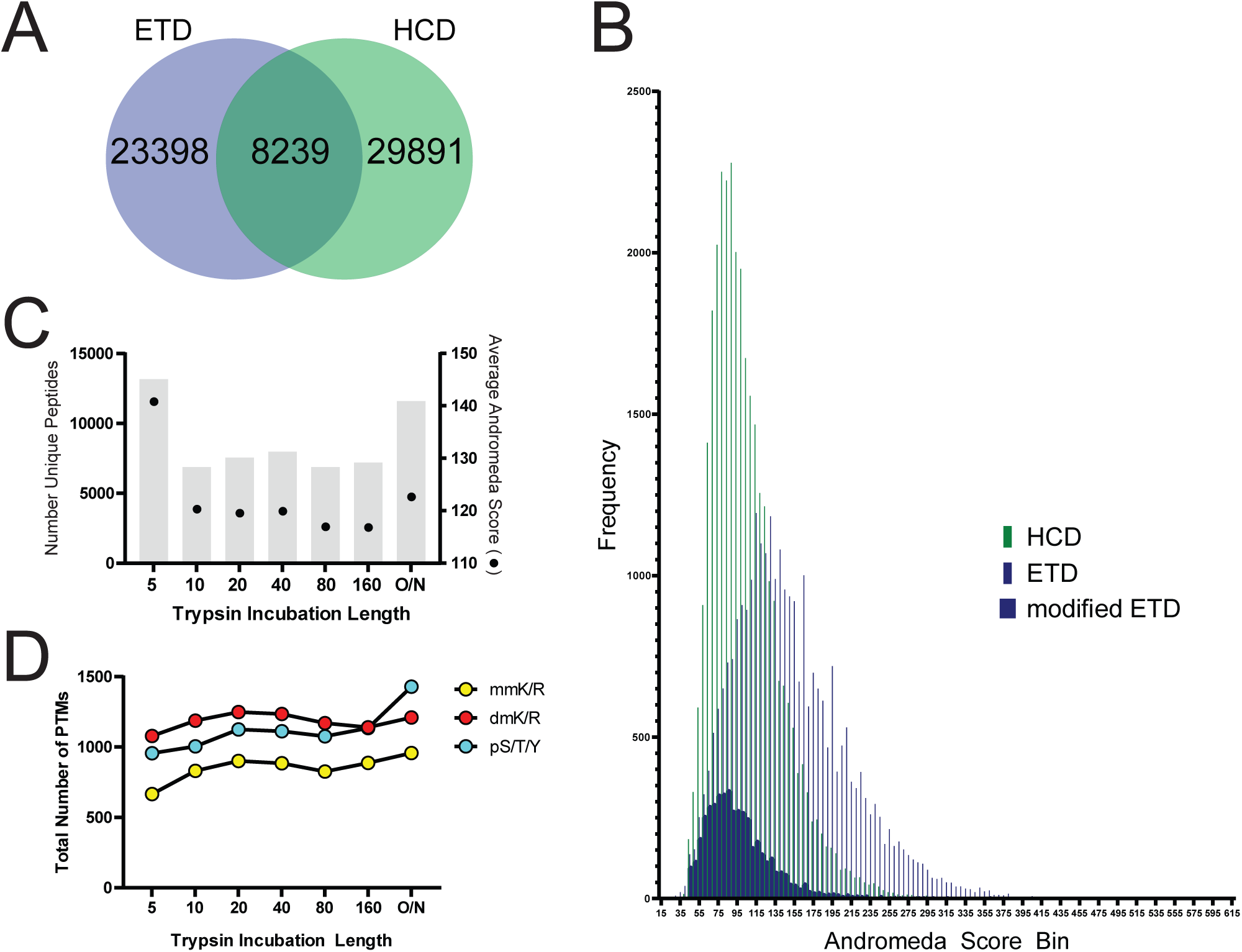
Middle-down ETD analysis of nucleoplasm contributes to greater coverage of Arg-rich RNA-binding proteins. ***(A)*** Venn diagram showing overlap between HCD and ETD fragmentation strategies of 61,528 unique peptides sequenced from nucleoplasm extract. ***(B)*** Andromeda scores of peptides fragmented by HCD (*thin green lines*) versus ETD (*thin blue lines*). Peptides with at least one PTM and sequenced by ETD are overlaid (*solid blue bars*). ***(C)*** The number of peptides sequenced (*grey bars*) and the average Andromeda score of a peptide (*black points*) at each trypsin incubation time length. ***(D)*** The total number of PTM across increasing trypsin incubation time lengths (monomethylation of Lysine/Arginine= mmK/R, *yellow*; dimethylation of Lysine/Arginine= dmK/R, *red*; phosphorylation of Serine/Threonine/Tyrosine= pS/T/Y, *turquoise*).

We then sought to assess the overall quality of our peptide spectral matches after expanding the search space of our raw spectral data using a non-standard targeted search accounting for increased missed cleavage events produced as a result of the shorter trypsin digestion times necessary for analyses of Arg-rich proteins (Supplemental Fig. S1D). Peptides at shorter trypsin digestion time lengths were high-confidence identifications with Andromeda scores comparable to the standard overnight digestion sample (Fig.4C). Interestingly, the 5-minute digestion produced the most unique peptides of any time point, and averaged higher Andromeda scores (140.78) than the standard overnight digestion sample (122.63) (Fig.4C).

When analyzing peptides modified by methylation or phosphorylation, the reliance on shorter trypsin incubation lengths and ETD fragmentation strategies becomes even more apparent. The majority of peptides with at least one PTM are successfully matched after ETD fragmentation across all limited trypsin proteolysis time lengths tested, only surpassed by HCD fragmentation in the standard overnight digestion sample, where peptides average 0.58 missed-cleavage events (Supplemental Fig. S1B, S1D). These modified peptides fragmented by ETD do not suffer from significantly reduced confidence scores, however, as they generally overlap with the Andromeda Scores of HCD-fragmented peptides (Fig.4B). As methylation of arginine or lysine does not affect the charge of the residue, Arg-rich peptides may thus be preferentially identified according to the data dependent HCD/ETD decision tree, at earlier digestion time points by ETD (Fig.4D, Supplemental Fig. S1B) [28]. Indeed, 764 out of 1076 peptides to Arg-rich proteins (71%) were fragmented by ETD (Supplemental Fig. S2D). Therefore, it is apparent that ETD fragmentation and limited digestion strategies enhance of the identification of Arg-rich sequences.

### BAD and RS motif algorithm resolves RNA-binding protein subgroups with distinct biological properties

RNA-binding proteins with BAD and RS domains have multiple binding partners that can change due to cellular condition, hypothesized to be regulated by PTMs [3, 57, 58]. To study the characteristics of two divergent yet similar Arg-rich protein subgroups, we implemented a scoring algorithm to select for peptides with alternating basic-acidic “BAD” or arginine-serine “RS” dipeptides normalized to peptide length, and focused the following analyses for peptides that scored above a stringent cutoff score (Supplemental Fig. S2A-B). The BAD algorithm selected 543 peptides, approximately 0.89% of all unique peptides sequenced (Supplemental Fig. S2A). These peptides matched to 237 proteins (Fig.5A, **Supplemental Table S3**). The RS algorithm selected 533 peptides (0.89%), identifying 117 RS proteins (Supplemental Fig. S2B, Fig. 5A, **Supplemental Table S4**). Although each requires several missed cleavages to be sequenced, BAD and RS proteins show similar Andromeda Scores to nonBAD/RS peptides (Supplemental Fig. S2C). Out of 354 total BAD and RS proteins identified by our algorithms, 45 proteins were members of both groups (Fig.5B). Based on previous studies to define U1-70K binding partners [3], 33 of these 45 shared proteins interact with U1-70K (∼73%) (Fig.5B), not including U1-70K itself. Many of these proteins are RBPs that are involved in RNA processing and exhibit elevated insolubility in AD [3, 59, 60].

**Figure 5.**
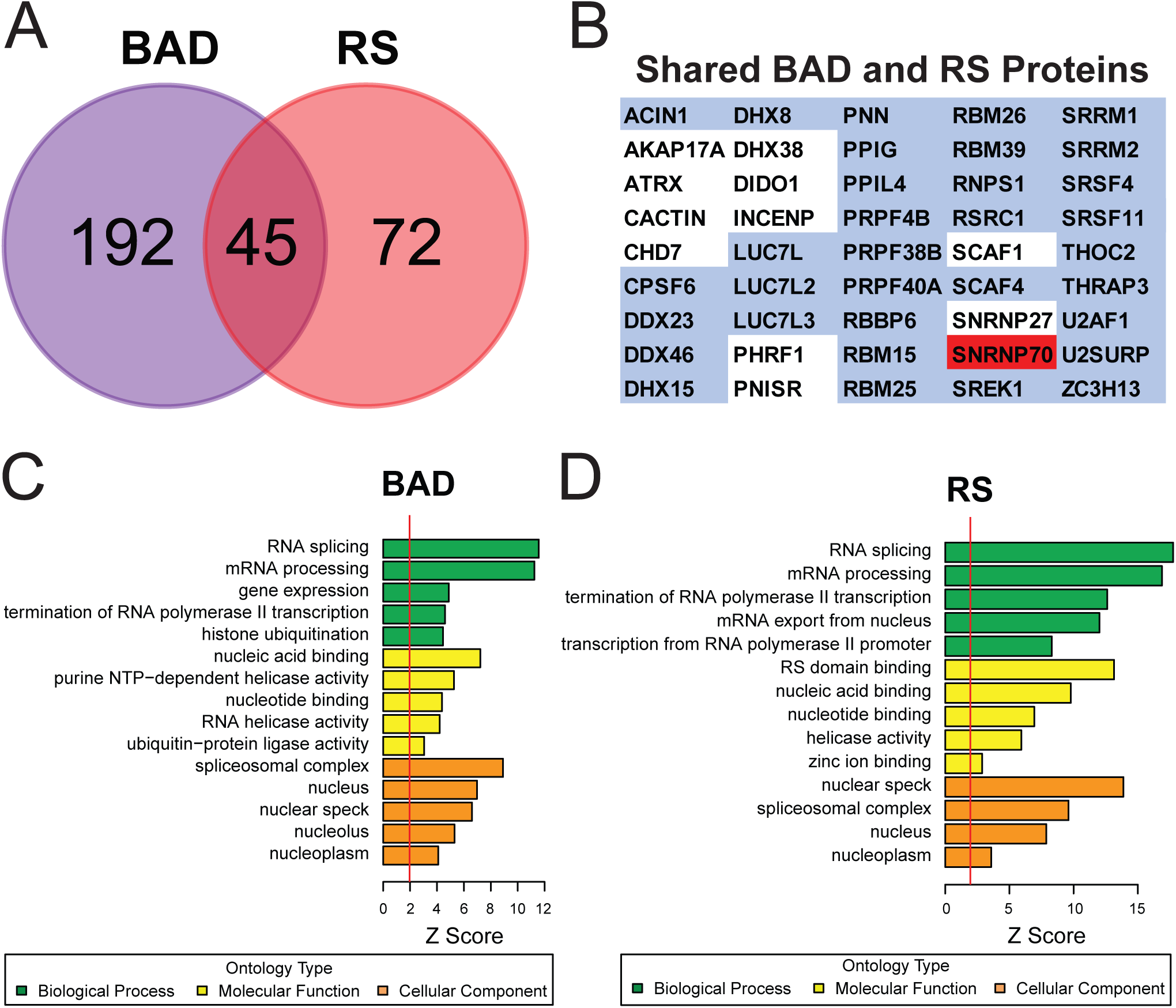
BAD and RS motifs resolve RNA-binding protein subgroups with distinct biological properties. ***(A)*** A scoring algorithm was developed (see methods) to select unique peptides that contain BAD- and RS-patterned primary sequences. ***(B)*** Of the 309 total proteins selected by the BAD and RS algorithms, 45 proteins had peptide matches that were considered both ‘BAD’ and ‘RS’, of which includes U1-70K (*red*). Of these 45, 33 were previously shown to interact with U1-70K (*blue* gene symbols). ***(C and D)*** Gene ontology (GO) analysis of BAD and RS proteins identified by our algorithm. Significant over-representation of an ontology term is reflected with a Z score greater than 1.96, equivalent to p<0.05 (to the right of the red line). ***(C)*** The BAD proteins (237) have the expected biological processes (RNA splicing and mRNA processing), and subcellular localization (spliceosomal complex and nucleus). ***(D)*** RS proteins (117) retain similar processes and localization (nuclear speckles and spliceosomal complex).

To examine the biological function of proteins containing BAD or RS domains, GO-elite analysis was performed, as compared to that of the total nucleoplasmic proteome identified by our analysis (Fig.5C-D). Using the 3,900 total nucleoplasmic gene symbols identified in this study as background, both the BAD and RS proteins are enriched in ‘RNA splicing’ and ‘mRNA processing’ functions. Both the BAD and RS proteins, in particular, are enriched with helicase activity (Fig.5C-D). Helicases remodel snRNA structure, which is a critical step in spliceosome re-arrangement, and various steps of RNA processing [61–66]. The RS proteins meanwhile, enrich for RS domain binding, as expected (Fig.5D). Furthermore, these algorithm-selected groups naturally partition to the correct subcellular locations, including ‘nuclear speckles’, the ‘spliceosomal complex’ and the ‘nucleoplasm’ (Fig.5C-D). Additionally, functions tangential to canonical splicing regulatory roles were parsed out by GO analysis, as there are many BAD proteins that ubiquitinate histones, including E3 ligases (Fig.5C) [67], while RS proteins aid in RNAPII transcription enhancement as well as mRNA export (Fig.5D). Thus, by utilizing ETD and bioinformatics approaches we were able to infer the biological function of two similar, yet distinct Arg-rich domains.

### Novel sites of phosphorylation and methylation on RBPs in the nuclear proteome revealed by middle-down ETD MS

Due to the increased proteomic coverage yielded by utilizing a combination of limited trypsin digestion and ETD fragmentation, we hypothesized that our method would achieve coverage of previously uncovered regions of the proteome, and in particular, BAD and RS proteins in the nucleoplasm fraction. All nucleoplasm peptides were then searched against all previously annotated sequences on the peptideatlas.org mass spectrometry repository database to determine if our sequences represented previously observed, or rather, unreported proteomic coverage [44]. A total of 29,114 residues mapping to the top isoform of each gene that were previously unobserved by mass spectrometry approaches, were uncovered by our middle-down ETD examination (Supplemental Fig.S3A). If this analysis is applied to all known alternate isoforms, we achieved coverage of a total of 101,438 residues, currently unreported on peptideatlas.org. The BAD and RS peptides alone matched to 2,410 and 3,096 previously un-sequenced residues mapping to the most common isoform, previously missed by conventional bottom-up MS methods (Supplemental Fig.S3A). With increased proteomic coverage, we hypothesized that a significant amount of unreported PTMs would also be identified. We thus compared PTM sites identified in this experiment to that of all PTMs previously identified by mass spectrometry and uploaded on phosphosite.org [42], which allows us to catalog unreported PTM sites identified in this study. A total of 681 unreported PTMs were identified in this experiment, with over half consisting of dimethylated arginine and phosphorylated serine (Supplemental Fig.S3B).

While only accounting for ∼1.5% of all residues detected (**Supplemental Table S2**), BAD and RS peptides consisted of ∼20% of all unreported residue coverage (Supplemental Fig.S3B). For example, peptides mapped to nucleoplasm-isolated U1-70K contributed to the coverage of 111 unreported residues, increasing total MS observation of U1-70K from ∼70% (Fig.1A) to 95% (Supplemental Fig. S4C). Importantly using middle-down ETD approaches within the complex nucleoplasmic sample mixture, we achieved near complete coverage (96%) of the LC1/BAD domain in endogenous U1-70K (Supplemental Fig. S4C).

### Arg-rich domains in RBPs contain combinatorial PTMs

As the Arg-rich BAD and RS domains have a high density of modifiable residues (Lys/Arg/Ser/Thr/Tyr), we hypothesized these domains may have an increased frequency of multiply-modified peptides. Indeed, the majority of BAD and RS peptides contained two or more PTMs, while less than 10% of nonBAD/RS peptides were multiply-modified (Fig.6A). While the majority of BAD and RS peptides were modified, nonBAD/RS peptides were overwhelmingly unmodified (89%) (Fig.6B). RS peptides, in particular, more frequently contained the maximum amount of PTMs (5) on a single peptide, than one or no PTMs at all (Fig.6B).

**Figure 6.**
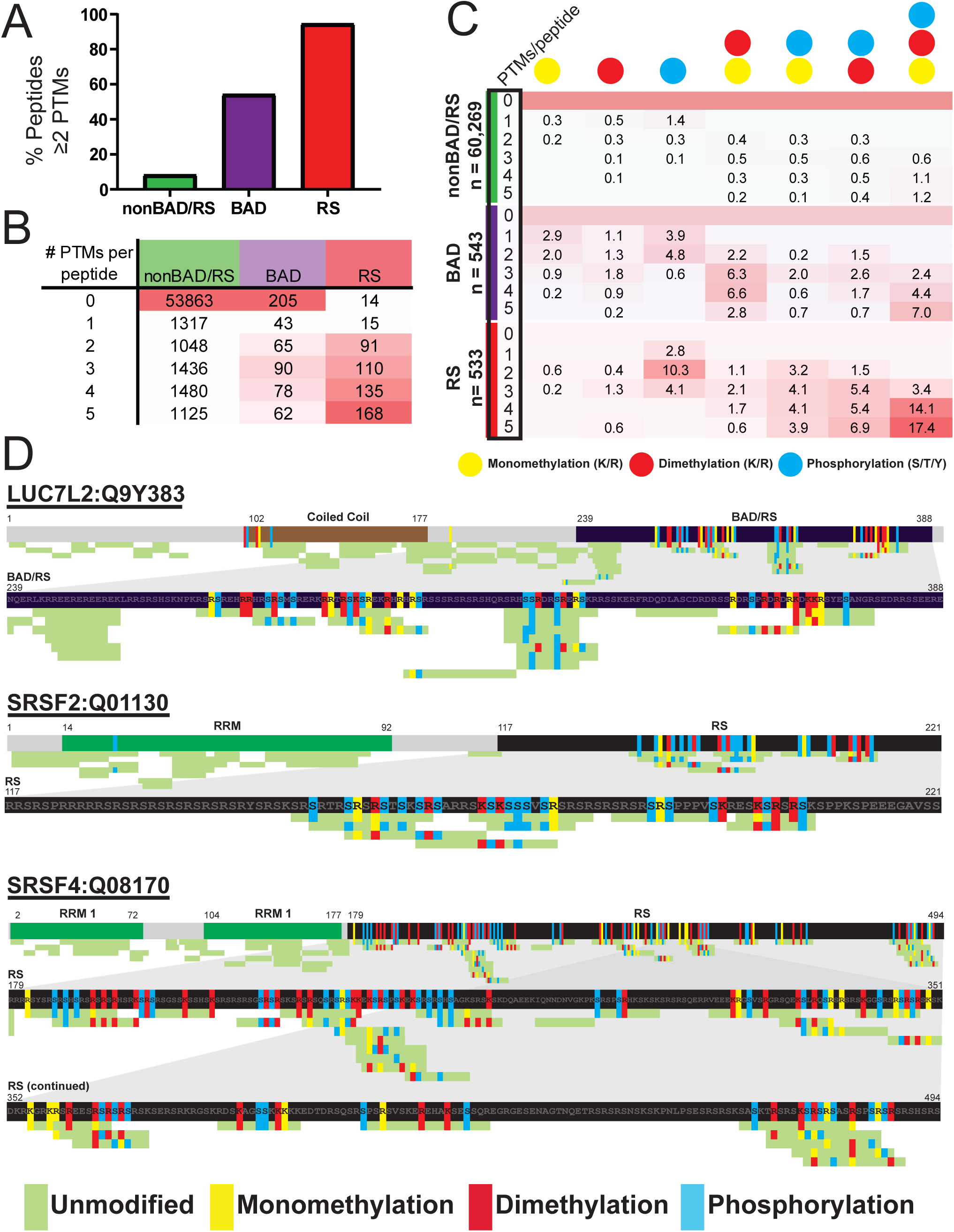
Arginine-rich RNA-binding proteins are enriched for PTMs. ***(A)*** The percentage of peptides containing two or more PTMs was calculated for nonBAD/RS, BAD and RS peptides. ***(B)*** The number of peptides containing no (0) to increasingly-modified states of PTM (1-5) were tallied and compared, using a heat map to illustrate occupancies of each subgroup. ***(C)*** The percentage of peptides containing uniform to double- or triple-combinations of PTM subtypes were calculated and compared between peptide groups (nonBAD/RS vs BAD vs RS), using a heat map to illustrate the frequency of each (monomethylation of Arginine/Lysine (*yellow*), dimethylation of Arginine/Lysine (*red*) or phosphorylated of Serine/Threonine/Tyrosine (*turquoise*). ***(D)*** Unique peptides were mapped to arginine-rich proteins LUC7L2, SRSF2, and SRSF4. Peptides are colored according to whether each residue is unmodified (*green*), monomethylated at an Arginine/Lysine (*yellow*), dimethylated at Arginine/Lysine (*red*) or phosphorylated at Serine/Threonine/Tyrosine (*turquoise*). High modification density was observed in the BAD and/or RS low-complexity (LC) domains of each of the three proteins. Sequences used to build this illustration are listed in **Supplemental Table S5**.

We next sought to determine whether BAD and RS peptides contained a combination of several PTM subtypes at once. Peptides belonging to nonBAD/RS, BAD and RS subgroups were classified according to the number of PTMs contained (0-5). The percentage of peptides belonging to uniform PTM states (mono-methylation, di-methylation, phosphorylation), double-PTM states (mono- and di-methylation, monomethylation and phosphorylation, dimethylation and phosphorylation) or a triple-PTM state (mono- and di-methylation and phosphorylation) were then calculated, and percentages were displayed as a heat map (Fig.6C). The majority of nonBAD/RS peptides were unmodified, although the most frequent PTM state was a single phosphorylation PTM. BAD peptides were enriched in combinatorial PTMs, with over 40% containing a combination of PTM subtypes on a single peptide (Fig.6C). RS domains were more frequently modified, as almost 3 out of every 4 peptides were modified by a combination of PTM subtypes (Fig.6C). Namely, ∼30% of all RS peptides contained all three PTM subtypes searched for in a single peptide (Fig.6C).

This was further illustrated when we performed peptide mapping within several highly-rated BAD and RS proteins, including LUC7L2, SRSF2 and SRSF4. Indeed, the Arg-rich BAD/RS domains were uniquely enriched in methylation and phosphorylation (Fig.6D**, Supplemental Table S5**). Compared to peptides mapping to surrounding domains, BAD and RS domains have a complex combinatorial signature of PTM. As PTMs are an essential regulator of the structure and function of RBPs [68–72], we next sought to further characterize the frequency of these PTMs in the nucleoplasm proteome.

### Arg-rich domains in RBPs are densely modified

We next performed PTM quantification on BAD and RS peptides, as compared to the background nucleoplasm proteome. For instance, we calculated the average frequency of a PTM within a peptide, divided by the number of potential modifiable (Lys/Arg/Ser/Thr/Tyr) residues within a peptide. On average, approximately 22% of Lys/Arg/Ser/Thr/Tyr residues within the BAD peptides are modified (Fig.7A). This is in stark contrast to ∼4.5% of residues modified in the remainder of the nucleoplasm (nonBAD/RS) proteome sequenced (Fig.7A). The RS peptides are similarly increased in methylation and phosphorylation, as nearly a quarter (23%) of Lys/Arg/Ser/Thr/Tyr residues within RS peptides are modified (Fig.7A). In sum, the BAD and RS peptides are approximately five-fold more likely to contain lysine methylation, arginine methylation or serine phosphorylation, compared with nonBAD/RS sequences identified in the nucleoplasm (Fig.7A).

**Figure 7.**
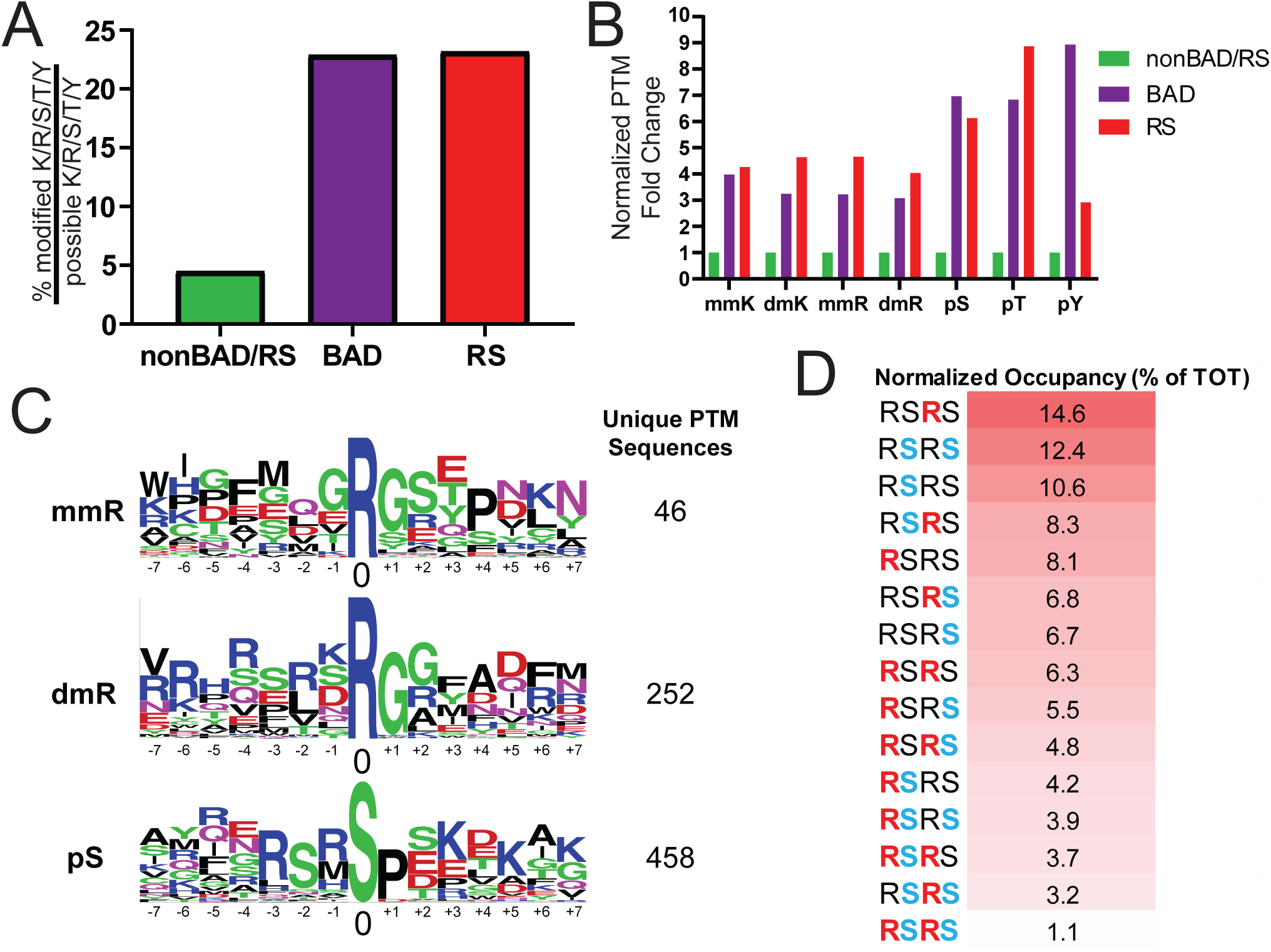
Arginine methylation and serine phosphorylation are enriched in RSRS motifs. ***(A)*** The density of modified residues (K/R monomethylation, K/R dimethylation, S/T/Y phosphorylation) within a peptide divided by the density of the total residues (K/R/S/T/Y) within the same peptide was averaged for peptide subgroups (nonBAD/RS, BAD and RS) and presented as an averaged percentage. Approximately 22% of BAD peptide residues are post-translationally modified, compared to 4.5% of background nucleoplasmic proteome. In comparison, approximately 23% of all residues in RS peptides are post-translationally modified. ***(B)*** Fold enrichment of PTM frequency for lysine, arginine, and serine were observed in BAD and RS peptides, normalized to the PTM frequency observed in nonBAD/RS peptides. ***(C)*** Motif Logos illustrating residue frequencies adjacent to a center modified residue was performed for monomethylated arginine (mmR), dimethylated arginine (dmR) and phospho-serine (pS). ***(D)*** Fifteen combinations of arginine methylation and serine phosphorylation observed within RSRS sequences were normalized compared by a percent of total occurrences calculation and a heat map illustration.

Next, we sought to determine if the PTM density increase observed is due purely to an increased number of modifiable residues or rather, a true increase in PTM frequency in BAD and RS regions compared to residues in nonBAD/RS regions. We divided the PTM frequencies observed in BAD and RS peptides by the frequencies observed in the background nucleoplasm proteome (nonBAD/RS) to estimate the relative fold increase in PTM frequency. We found markedly increased methylation and phosphorylation fold changes, indicating that the hyper-modification observed within BAD and RS sequences is not simply caused by an increase in modifiable residue density within these regions, but also a naturally increased likelihood for residues in these domains to be modified (Fig.7B). For example, serine in particular is almost six times more likely to be phosphorylated in BAD and RS domains, compared to the rest of the proteome, even when normalizing for the presence of serine by our selection criteria (Fig.7B). Thus, by increasing proteomic coverage with middle-down ETD, we discovered that Arg-rich domains contain markedly increased PTM densities.

### Phosphorylation and methylation favor RSRS motifs, yet do not frequently co-occur in this motif

Due to the high level of combinatorial PTMs detected within Arg-rich low complexity sequences, we analyzed PTM motif sequences in the nucleoplasm sample. To determine if increased proteomic coverage influences phosphorylation and methylation PTM motif sites, a motif analysis was performed to determine if consensus sequences were observed at sites of arginine methylation and serine phosphorylation PTMs. Our nucleoplasm middle-down ETD MS PTM-containing sequences were searched in 31-residue width windows using the Motif Logo tool offered by phosphosite.org to detect increased frequencies of residues adjacent to a central modified residue for monomethylated arginine (mmR), dimethylated arginine (dmR) and phosphorylated serine (pS) (Fig.7C) [42]. Importantly, the PTM motif searches were conducted in isolation, without consideration of other PTM subtypes. As expected, the canonical RGG/GAR (glycine-arginine-rich) motif was found as the favored consensus motif for arginine monomethylation [73–75]. Unexpectedly, an “RSRS” motif featured prominently for not only serine phosphorylation [76], but also arginine dimethylation (Fig.7C). The difference of motifs between mono-methylation and di-methylation modification states of arginine highlights the utility of the middle-down ETD strategy and alternate MS approaches in uncovering unreported features of the proteome.

To assess whether serine phosphorylation and arginine methylation co-occur within RSRS motifs, a search was performed to select peptide sequences containing RSRS motifs. Interestingly, although arginine dimethylation and serine phosphorylation occur most frequently within BAD and RS peptides in nucleoplasmic RNA-binding proteins, these PTMs do not tend to co-occur within an RSRS motif. Namely, the most frequent modification states of the RSRS motif are either the central arginine residue(s) methylated (14.6%) or double serine phosphorylation (12.4%) (Fig.7D). This disparity of modification states suggests a complex layer of co-regulatory cross-talk between arginine and serine residues within Arg-rich domains.

## DISCUSSION

Mass spectrometry has achieved great advances towards the goal of globally characterizing PTMs within proteins [77], yet significant gaps in understanding remain. Here we describe a method to examine the PTM diversity of Arg-rich domains, re-purposing a technique first conceptualized by others to explore the biology of histone tails [16, 24, 32, 78]. The power of middle-down ETD is that this method can map unique profiles of combinatorial PTMs within Arg-rich proteins, many of which are RBPs that aggregate in neurodegenerative disease [3, 36]. This approach could be extended to diseased tissues to understand how PTM status changes and correlates with disease state or progression. This technique can be further used to determine the PTM profiles of Arg-rich domains across the proteome, eventually illuminating key regulatory steps in RNA processing and metabolism.

Here, we achieved full proteomic sequence coverage of the recombinant arginine-rich U1-70K LC1/BAD domain by a combination of limited proteolysis and ETD approaches. We then expanded the approach for the proteomic analysis of nucleoplasm proteins isolated from mammalian cells, many of which are Arg-rich RBPs. The use of ETD sequencing on arginine-rich proteins contributed to the coverage of thousands of residues currently un-reported on peptideatlas.org [44, 45], and hundreds of PTMs previously unannotated on phosphosite.org [42]. Future examination with alternative proteolytic enzymes, advanced search engines and high resolution Orbitrap MS/MS strategies will greatly aid in the advancement of unreported combinatorial PTM identification and quantification.

A vital discovery achieved by our analysis is the combinatorial nature of PTMs within Arg-rich domains, highlighted by a shared RSRS motif between arginine dimethylation and serine phosphorylation in nucleoplasm proteins [73, 79]. Serine-arginine protein kinases (SRPKs) and protein arginine methyltransferase (PRMT) enzymes modify arginine-rich domains in RNA-binding proteins [12, 68, 80, 81]. Due to the proximal relationship of methylation and phosphorylation PTMs, cross-talk between adjacent arginine and serine residues within RSRS motifs is likely. Phosphorylation and methylation have been previously shown to functionally oppose one another. For example, arginine methylation at RxRxxS/T motifs, similar to RSRS motifs, have emerged as an important negative regulator of serine phosphorylation. The mouse homolog of forkhead box O1 (FOXO1) is methylated in vitro and in vivo at its Akt recognition motif site at R248 and R250, blocking phosphorylation of nearby S253 and preventing FOXO1 nuclear export [82]. Conversely, phosphorylation also blocks methylation in certain contexts. Namely, phosphorylation of RNA polymerase II (RNAPII) at its carboxy terminal domain prevented symmetric arginine dimethylation at R1810, integral to SMN protein interaction [83, 84]. It is evident that there is functional cis-acting crosstalk between site-specific methylation and phosphorylation PTMs that is context-dependent. Therefore, the overall ratio between methylation and phosphorylation in arginine-rich domains may critically define the protein function. The global identification of novel sites and further interrogation of the crosstalk between methylation and phosphorylation of RS motifs is warranted and may be revolutionized by this type of mass spectrometry approach.

The appearance of a range of arginine methylation states reflect a dynamic PTM process, in which protein function may be tuned by varying degrees of methylation of contiguous arginines. Interestingly, PRMTs appear to have a broad yet essential role in regulating alternative splicing. Type II PRMT5 was recently shown to be preferential substrates of RNA-binding proteins. PRMT5-depleted cells critically trigger changes in gene expression, cell-cycle de-regulation and alternative splicing [81, 85]. Another type II enzyme, PRMT9, regulates alternative splicing by methylation of spliceosome-associated protein 145 (SAP145) at R508, priming SAP145 for interaction with the protein SMN [86]. Thus, arginine methylation is likely an essential regulator of alternative splicing, and the identification residue-specific of arginine methylation sites in splicing RBPs can reveal novel and essential insights to structural and functional inquiry of these proteins. In the future, performing ETD MS on PRMT inhibitor-treated cells may reveal site-specific targets to individual residues within BAD and RS domains. This may offer insight to the therapeutic capacity of individual PRMT enzyme inhibition in a disease-specific basis. Numerous lines of evidence suggest that further characterization of RBP PTM status will increase our understanding of multiple steps in RNA processing [7, 9, 12, 87].

A fundamental observation made in this study is that Arg-rich domains are densely modified by PTMs. These modifications likely regulate numerous aspects of protein function and localization. Furthermore, the middle-down ETD approach described here can be modified to study similarly Arg-rich dipeptide repeats, including poly-GR and poly-PR dipeptides repeats, translated from an intronic expansion repeat in *C9ORF72* of ALS. These large polypeptide expansion repeats may be sinks of arginine methylation, disrupting normal PTM status of arginine-rich RBPs in ALS [88]. This sequencing technology could allow for the mapping of site-specific PTM within these repeats, as well as a first opportunity to relatively quantitate DPRs of various lengths above disease-thresholds from patient samples. At the moment, antibodies are used to confirm DPR presence [89, 90], which do not to resolve exact DPR length.

One limitation of our study is the lack of PTM enrichment strategies, including IMAC and PTM immunoaffinity approaches, limiting the true depth of our analysis of the nucleoplasmic proteome. The intriguing possibility that one PTM-enriched sample may purify peptides with combinatorial PTMs is intriguing, and new, powerful search engines and algorithms are currently being developed to handle the expanded search space of multiple PTMs, such as MSFragger and TagGraph [91, 92]. However, by cellular fractionation methods we were able to capture and sequence combinatorial-modified peptides after a two-step database search using a 1% PSM and protein FDR cutoff [40].

Importantly, the unique pattern of ETD fragmentation itself aids in the sequencing of modified peptides [33]. The generation of many c-and z-type fragment ions contributes to the generation of unique fragment ions across a wide range of m/z values, increasing the likelihood of accurate peak calling by search engine algorithms, compared to HCD/CID fragmentation platforms. We performed our analyses on a Orbitrap mass spectrometer platforms, with a high-resolution precursor (MS1) scan and a low-resolution ion-trap MS/MS scan. The CID MS/MS scan post-ETD fragmentation suffers from a lack of resolution needed to identify PTMs from others with similar monoisotopic mass changes (e.g. deamidation and citrullination, lysine acetylation and tri-methylation). Lysine acetylation (42.011 ± 0.004 Da) and tri-methylation (42.047 ± 0.002 Da) variable modifications were excluded in this current study, as the change in mass between the divergent PTM fragment ions cannot be differentiated using a low-resolution ion-trap MS/MS scan (product ion mass tolerance = 0.6 Da). Future studies resolving these biologically-significant PTMs is warranted, and may be achieved using an high-resolution MS/MS scan.

Notably, a number of orthogonal proteases may be used to plausibly increase and cross-validate sequence coverage of Arg-rich domains, including proteases that cleave non-specifically (Elastase), at acidic residues (AspN, GluC), at basic residues (ArgC, LysC) and at aromatic residues (Chymotrypsin). Multi-protease strategies such as Confetti have successfully increased coverage of the proteome [93]. For proteases that cleave at basic and acidic residues (ArgC, LysC, AspN, GluC), however, a limited trypsin proteolysis strategy must be utilized in a manner similar to that described in this study. Furthermore, cleaving U1-70K with the protease Chymotrypsin would yield an 81 residue peptide fragment containing the LC1/BAD domain. Mass spectrometry observation of the LC1/BAD domain within such a large peptide fragment from a complex mixture would be extremely unlikely, as would be other Arg-rich low complexity sequences. Although possibly providing complementary information, the use of proteases other than trypsin are not as amenable to the study of highly-basic low complexity Arg-rich domains from complex mixtures. Trypsin remains a robust protease that, while using middle-down strategies, generates highly-basic Arg-rich peptides of suitable lengths that are primed for ETD fragmentation and mass spectrometry analysis.

Finally, by illuminating previously “dark” regions of the Arg-rich proteome we have discovered densely-modified regions within specific classes of RBPs. This dataset is an immense resource for the RBP community that investigates the roles of PTMs in RNA processing. The site-specific PTMs identified herein may be leveraged in future mutagenesis studies in model systems to guide functional studies of these biologically-significant proteins. Increasing the catalog of PTMs could be key in understanding global regulation of the spliceosome and non-histone epigenetic influences on cellular biology, highlighting the ability of a novel mass spectrometry pipeline to discover previously unappreciated aspects of cell biology.

## Abbreviations

MS: mass spectrometry
ETD: electron transfer dissociation
PTM: post-translational modification
AD: Alzheimer’s disease
LC: liquid chromatography
RBP: RNA-binding proteins
PSM: peptide spectral match

## Acknowledgment

Support was provided by 5R01AG053960 and by the Accelerating Medicines Partnership for AD (U01AG046161). S.K. was supported by the Training Program in Biochemistry, Cell and Developmental Biology (T32GM008367) and Emory Neurology Department Training Fellowship (5T32NS007480-19). I.B. was supported by a pre-doctoral NINDS training grant (T32NS007480) and an individual NRSA grant (F31NS09385902). We thank Anita Corbett (Emory Department of Biology) for her helpful comments and suggestions.

## Supporting information

**The following supporting information is available free of charge at ACS website** http://pubs.acs.org

**Supplemental Table S1**: **List of U1-70K LC1/BAD peptides.** List of peptides mapping to U1-70K LC1/BAD.

**Supplemental Table S2. Nucleoplasmic middle-down ETD MS experiment summary statistics. S**ummed and averaged values of 61,283 total unique peptides, including the subgroups of nonBAD/RS (*grey*), BAD (*purple*) and RS (*red*) peptides.

**Supplemental Table S3. BAD protein and peptide list.** The 237 BAD proteins listed as gene symbols, with 543 peptides and accompanying BAD scores.

**Supplemental Table S4. RS protein and peptide list.** The 117 RS proteins listed as gene symbols, with 533 peptides and accompanying RS scores.

**Supplemental Table S5: List of LUC7L2, SRSF2, and SRSF4 peptides.** List of peptides mapping to LUC7L2, SRSF2 and SRSF4.

**Supplemental Figure S1:**
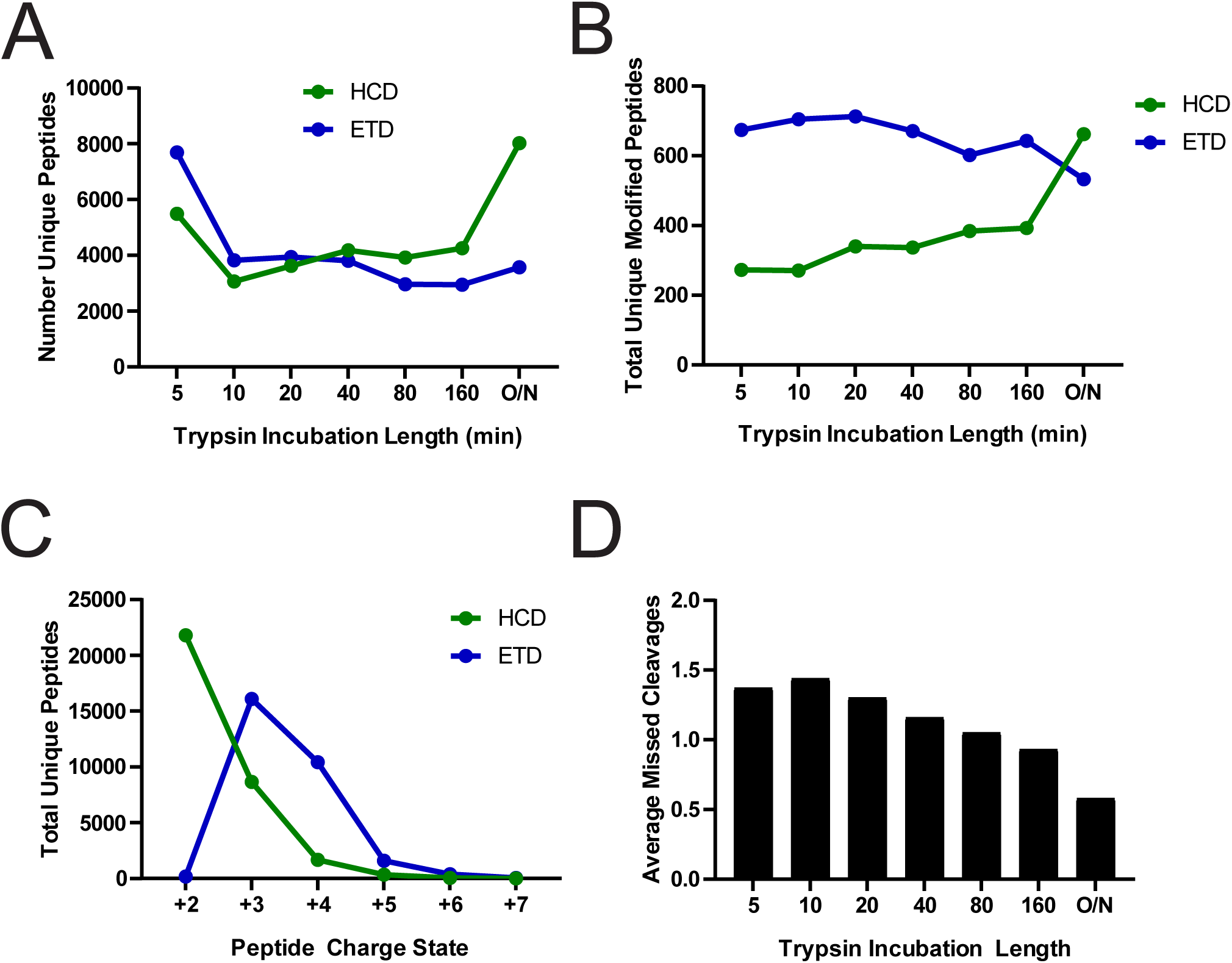
HCD and ETD fragmentation of nucleoplasmic sample enhances identification of unique peptides. ***(A)*** At early trypsin times, ETD MS/MS fragmentation produces the majority of sequenced peptides. HCD MS/MS is the most utilized fragmentation method from 40 minutes on. ***(B)*** The number of unique modified peptides sequenced using ETD outpaces HCD at all trypsin digest time points except for overnight (O/N) digestion. ***(C)*** As determined by the decision tree approach [28], peptides with charges greater than +2 are preferentially sequenced by ETD. ***(D)*** Average missed cleavages events decrease as trypsin incubation times increase.

**Supplemental Figure S2:**
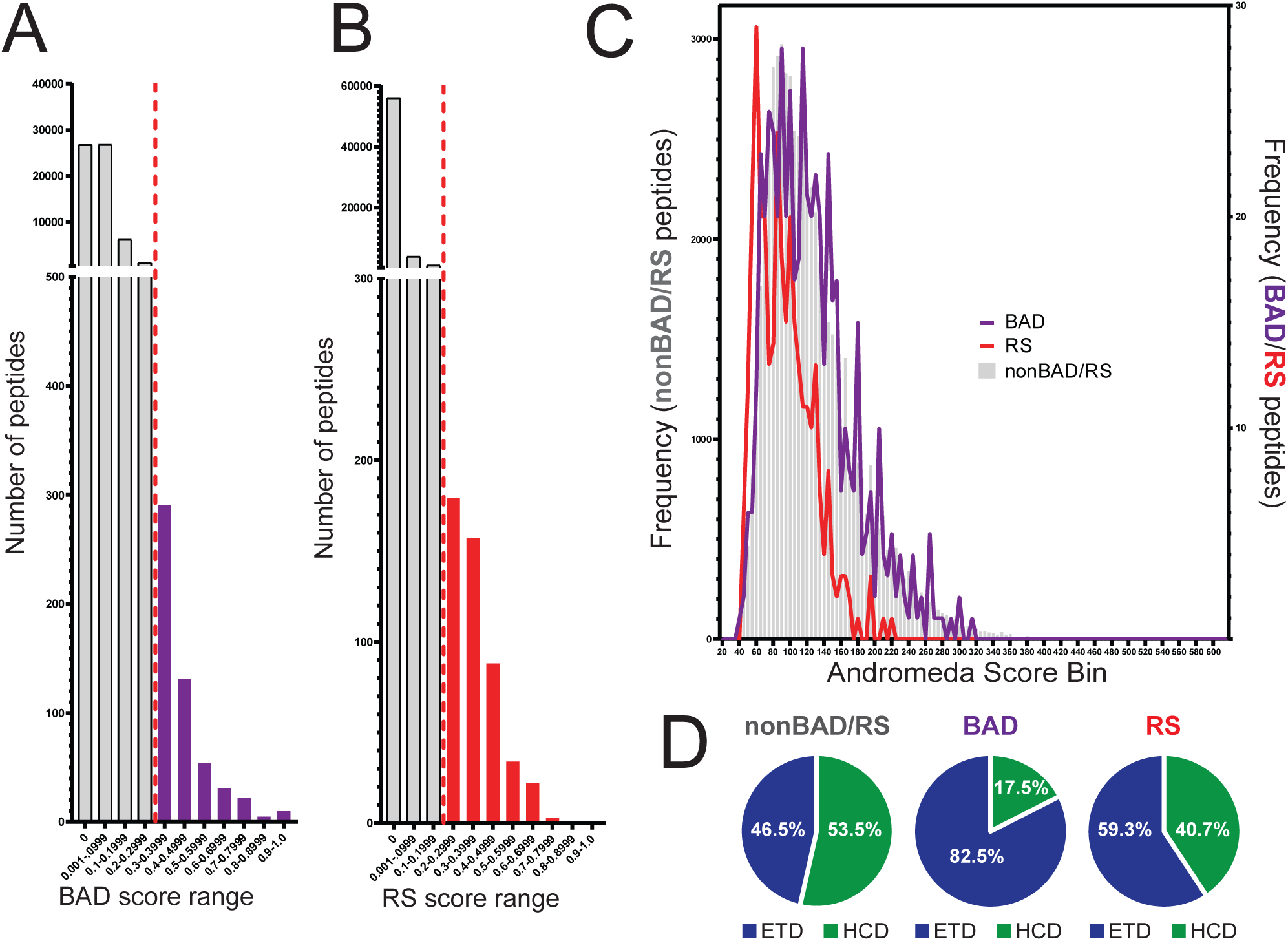
A BAD and RS motif algorithm stringently selects high-confidence peptides that are preferentially fragmented by ETD. **(A-B)** A scoring algorithm was developed (see methods) to select identified unique peptides that contain BAD-and RS-patterned primary sequence. A total of 61,283 unique peptides were scored using the scoring algorithms, and a conservative score cutoffs of 0.3 or greater (BAD) and 0.2 or greater (RS) were used for each algorithm. ***(A)*** A total of 543 peptides (237 proteins) were scored as BAD peptides, approximately 0.89% of all peptides sequenced. ***(B)*** A total of 533 peptides (117 proteins) were scored as RS peptides, approximately 0.87% of all peptides sequenced. ***(C)*** Histogram analysis of the MaxQUANT Andromeda Scores of nonBAD/RS (*grey*), BAD (*purple*) and RS (*red*) peptides. ***(D)*** Frequency of HCD (*green*) or ETD (*blue*) fragmentation strategies yielding successfully-identified unique peptides to nonBAD/RS, BAD or RS peptide subgroups.

**Supplemental Figure S3:**
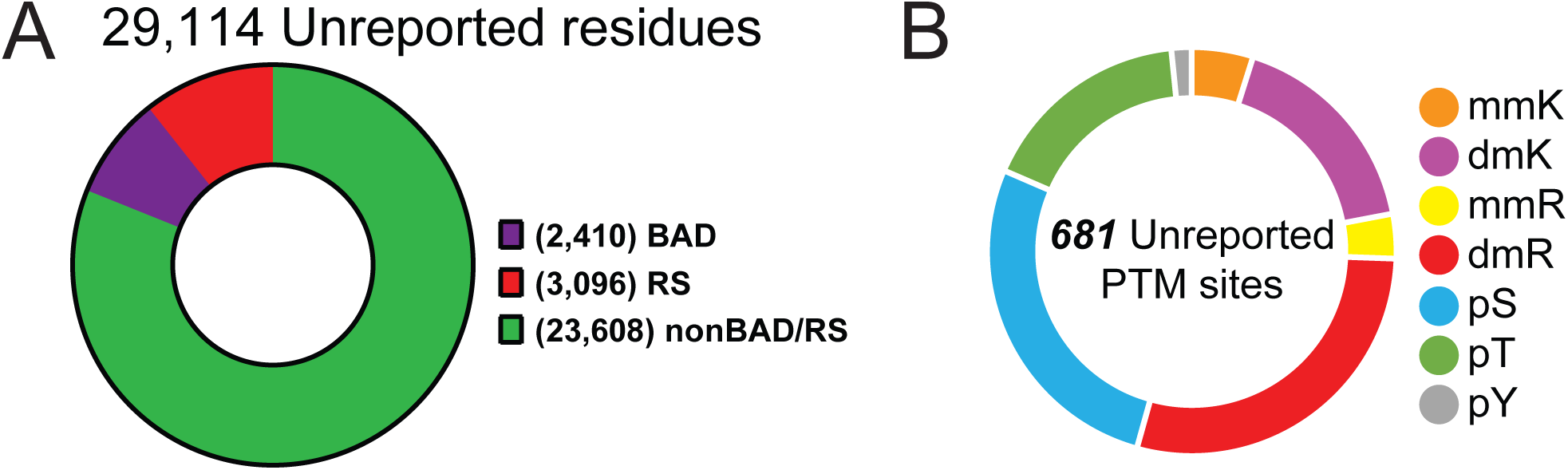
middle-down ETD approaches improve protein coverage and PTM identification. ***(A)*** A total of 29,114 novel residues unannotated on Peptide Atlas (http://www.peptideatlas.org/) were detected by this middle-down ETD MS approach. The BAD peptides totaled novel coverage of 2,410 residues, while RS peptides totaled 3,096 previously unmatched residues. ***(B)*** A total of 681 novel PTM sites were identified (mmR=monomethyl arginine, *yellow*; dmR=dimethyl arginine, *red*; mmK=monomethyl lysine, *orange*; dmK=dimethyl lysine, *magenta*; pS=phospho-serine, *turquoise*; pT=phospho-threonine, *green*; pY=phospho-tyrosine, *grey*).

**Supplemental Figure S4:**
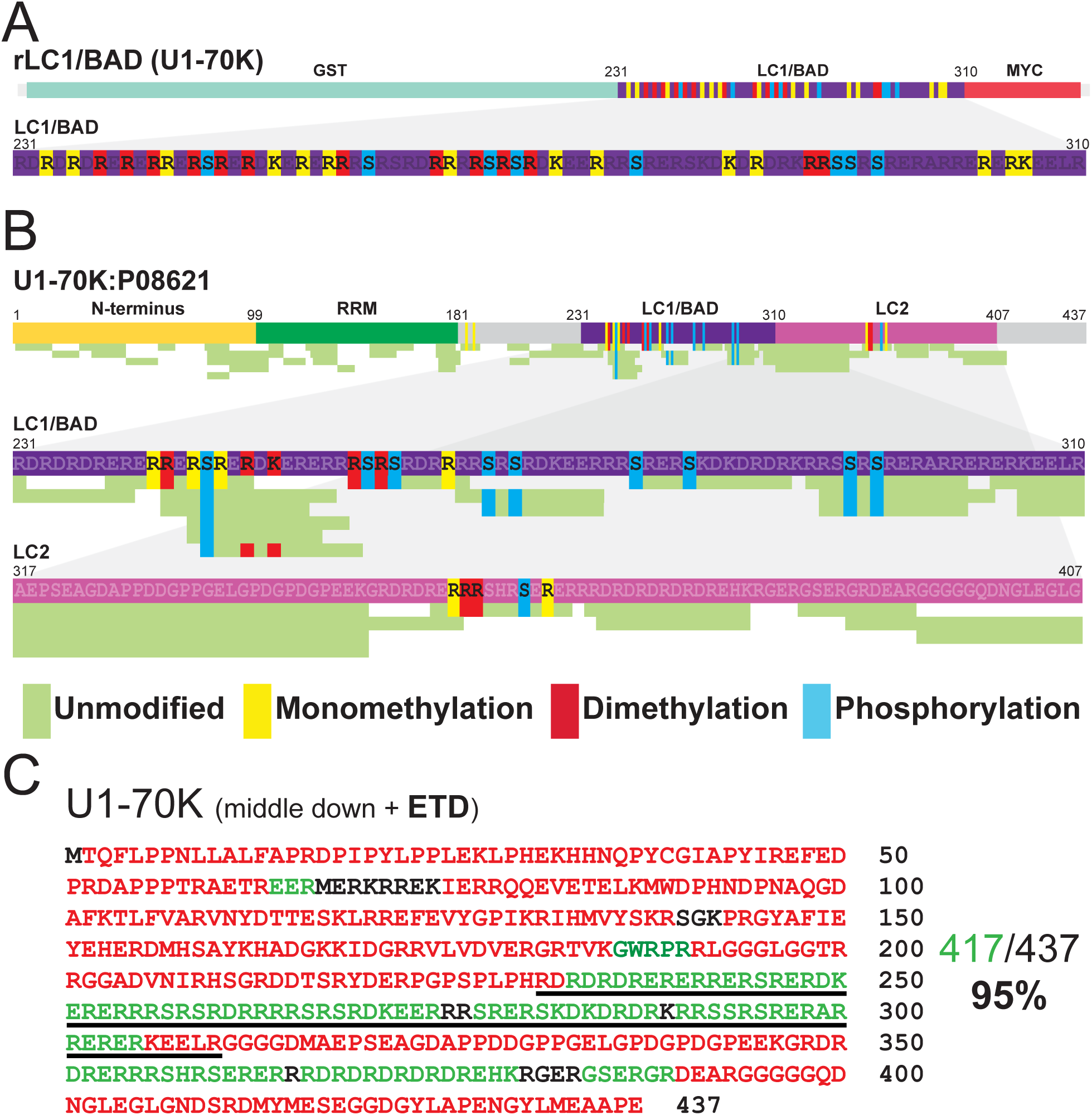
Middle-down ETD MS strategy provides novel sequence coverage of U1-70K arginine-rich low complexity domains. ***(A)*** Summation of modified peptide spectral matches (PSMs) of all trypsin digestion times and ETD/HCD dissociation methods on the LC1/BAD domain of the purified recombinant LC1/BAD protein. Peptides are colored according to whether each residue is unmodified (*green*), monomethylated at an Arginine/Lysine (*yellow*), dimethylated at Arginine/Lysine (*red*) or phosphorylated at Serine/Threonine/Tyrosine (*turquoise*). Using middle-down ETD, near complete coverage of the LC1/BAD domain was achieved, identifying 8 phosphorylation sites and 23 methylation sites (12 mono-methylated arginines, 13 di-methylated arginines, and four mono-methylated lysines) in total. ***(B)*** Amino acids 231-310 of the HEK293T-native U1-70K LC1/BAD domain was almost completely sequenced (76/80 residues, 95%), identifying 9 phosphorylation sites with 9 total methylation sites. By comparison, the U1-70K LC2 domain, without BAD or RS motifs but similar arginine content, contains less methylation and phosphorylation PTM. ***(C)*** Residues in red are those that have been previously observed by mass spectrometry analysis. U1-70K has 437 total amino acids, with 70% sequence coverage (306 AAs, *red*) currently deposited on the repository peptideatlas.org. As a result of our ETD analysis of nucleoplasm extract alone, we identified 111 new amino acids (*green*). This has led to an increase to 95% total sequence coverage of U1-70K by middle-down ETD MS approaches.

**Supplemental Figure S5:**
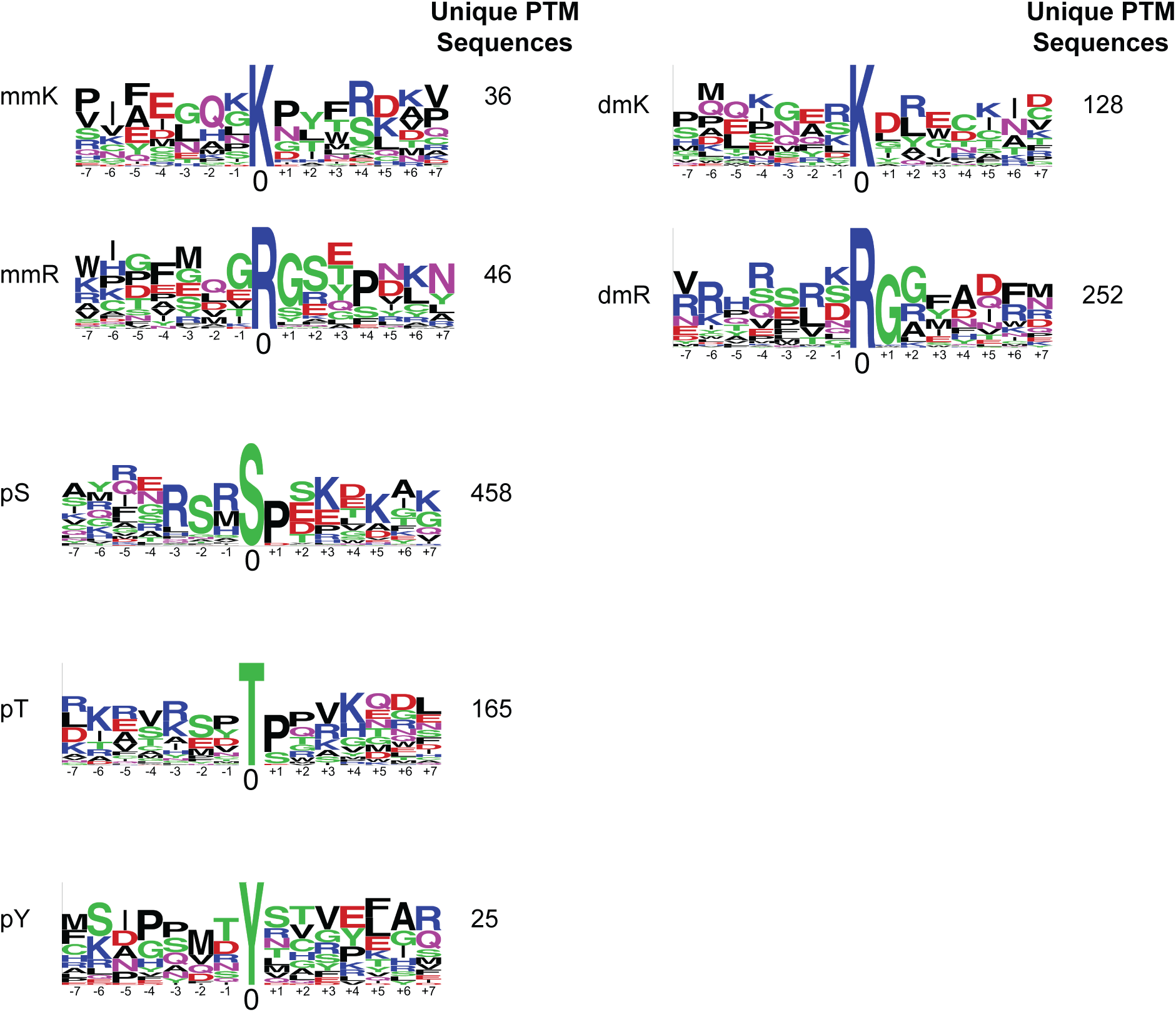
Motif analysis of nucleoplasmic PTM-modified peptide sequences. Unique PTM-containing sequences were searched using the Motif Logo tool on PhosphoSite.org to illustrate the frequencies of residues with increased occupancies at positions relative to a center modified residue (mmR=monomethyl arginine, dmR=dimethyl arginine, mmK=monomethyl lysine, dmK=dimethyl lysine, pS=phospho-serine, pT=phospho-threonine, pY=phospho-tyrosine)

